# Mitophagy Facilitates Cytosolic Proteostasis to Preserve Cardiac Function

**DOI:** 10.1101/2024.11.24.624947

**Authors:** David R. Rawnsley, Moydul Islam, Chen Zhao, Yasaman Kargar Gaz Kooh, Adelita Mendoza, Honora Navid, Minu Kumari, Xumin Guan, John T. Murphy, Jess Nigro, Attila Kovacs, Kartik Mani, Nathaniel Huebsch, Xiucui Ma, Abhinav Diwan

**Author notes:** indicated contributed equally (David R. Rawnsley, Moydul Islam, Chen Zhao). **Corresponding Author**: Abhinav Diwan, M.D., Professor of Medicine, Cell Biology and Physiology, Obstetrics and Gynecology; and Xiucui Ma, Ph.D., Assistant Professor of Medicine, Washington University School of Medicine, Division of Cardiology, 660 S. Euclid, CSRB-NTA 827, St. Louis, MO 63110.

## Abstract

**Background:** Protein quality control (PQC) is critical for maintaining sarcomere structure and function in cardiac myocytes, and mutations in PQC pathway proteins, such as CRYAB (arginine to glycine at position 120, R120G) and BAG3 (proline to lysine at position 209, P209L) induce protein aggregate pathology with cardiomyopathy in humans. Novel observations in yeast and mammalian cells demonstrate mitochondrial uptake of cytosolic protein aggregates. We hypothesized that mitochondrial uptake of cytosolic protein aggregates and their removal by mitophagy, a lysosomal degradative pathway essential for myocardial homeostasis, facilitates cytosolic protein quality control in cardiac myocytes.

**Methods:** Mice with inducible cardiac myocyte specific ablation of TRAF2 (TRAF2icKO), which impairs mitophagy, were assessed for protein aggregates with biochemical fractionation and super-resolution imaging in comparison to floxed controls. Induced pluripotent stem cell (iPSC)-derived cardiac myocytes with R120G knock-in to the *CRYAB* locus were assessed for localization of the CRYAB protein. Transgenic mice expressing R120G CRYAB protein (R120G-TG) were subjected to both TRAF2 gain-of-function (with AAV9-cardiac Troponin T promoter-driven TRAF2 transduction) and TRAF2 loss-of-function (with tamoxifen-inducible ablation of one *Traf2* allele) in cardiac myocytes to determine the effect of mitophagy modulation on cardiac structure, function, and protein aggregate pathology.

**Results:** Cardiomyocyte-specific ablation of TRAF2 results accumulation of mitochondrial and cytosolic protein aggregates and DESMIN mis-localization to protein aggregates. Isolated mitochondria take up cardiomyopathy-associated aggregate-prone cytosolic chaperone proteins, namely arginine to glycine (R120G) CRYAB mutant and proline to lysine (P209L) BAG3 mutant. R120G-CRYAB mutant protein increasingly localizes to mitochondria in human and mouse cardiomyocytes. R120G-TG mice demonstrate upregulation of TRAF2 in the mitochondrial fraction with increased mitophagy as compared with wild type. Adult-onset inducible haplo-insufficiency of TRAF2 resulted in accelerated mortality, impaired left ventricular systolic function and increased protein aggregates in R120G-TG mice as compared with controls. Conversely, AAV9-mediated TRAF2 transduction in R120G-TG mice reduced mortality and attenuated left ventricular systolic dysfunction, with reduced protein aggregates and restoration of normal localization of DESMIN, a cytosolic scaffolding protein chaperoned by CRYAB, as compared with control AAV9-GFP group.

**Conclusions:** TRAF2-mediated mitophagy in cardiac myocytes facilitates removal of cytosolic protein aggregates and can be stimulated to ameliorate proteotoxic cardiomyopathy.

## Introduction

Protein quality control (PQC) in cardiac myocytes is critical for maintaining normal sarcomere structure and function and ensuring cellular viability over a lifespan spanning many decades in humans. Cardiac myocytes possess intricate and overlapping protein quality control mechanisms to maintain proteins in their physiologic location and function.^1, 2^ It is therefore not surprising that mutations in PQC pathway proteins ^3–6^ are causally implicated in human cardiomyopathy and heart failure, often in an autosomal dominant fashion indicating a gain of toxic function. These mutations, while rare, result in accumulation of misfolded aggregate-prone proteins as protein aggregates in cardiac myocytes and cause proteotoxicity resulting in cardiac myocyte death and dysfunction.^7–9^ Understanding the mechanisms for how protein aggregates and aggregate-prone proteins are removed will spur development of novel targeted therapies to ameliorate protein aggregates and treat cardiomyopathies.

In physiology, misfolded and damaged proteins are targeted for removal by covalent ligation of an ubiquitin moiety in chains (i.e. poly-ubiquitination) that targets proteins for degradation via the ubiquitin-proteasome pathway.^10^ Autophagy, a lysosomal degradative pathway, constitutes a parallel PQC mechanism and sequesters poly-ubiquitinated proteins (as well as other cargo such as damaged organelles) within double-membrane bound autophagosomes, followed by fusion with lysosomes for intra-lysosomal degradation.^11^ Autophagy is upregulated in the setting of proteasome dysfunction or insufficiency, as a backup pathway to facilitate PQC.^12, 13^ However, accumulating evidence points to dysregulation of both the ubiquitin-proteasome machinery and the autophagy-lysosome pathway in cardiomyopathy, especially in the setting of protein aggregate pathology in the myocardium.^1, 11, 14–16^ A case in point that illustrates the critical requirement for cytosolic PQC is DESMIN, a sarcomere-associated protein that scaffolds the sarcomeres and mitochondria in parallel proximity for functional coupling. CRYAB, a small heat shock protein, is highly enriched in cardiac myocytes^17^ and chaperones DESMIN to its normal location. A missense mutation in the cognate gene *CRYAB* results in an arginine to glycine (R120G) change and causes cardiomyopathy and heart failure inherited in an autosomal dominant fashion.^7^ The R120G mutant CRYAB protein forms aggregates which sequester DESMIN and other client proteins, leading to their abnormal localization and functional deficiency. CRYAB is degraded by both the ubiquitin-proteasome system^18, 19^ and the autophagy-lysosome pathway.^16^ While strategies to stimulate these pathways are partially effective in attenuating cardiac proteotoxicity^16, 20–22^, it is not known if alternate cellular pathways exist and can be targeted to facilitate removal of aggregates and restore normal sarcomeres.

A recent exciting discovery^23^ uncovered mitochondrial import of aggregated cytosolic proteins in yeast. This pathway was termed MAGIC (<u>M</u>itochondria <u>A</u>s <u>G</u>uardians In <u>C</u>ytosol)^23, 24^ with the implication that mitochondrial uptake of misfolded or damaged proteins plays a role in cytosolic PQC in homeostasis and under stress, a premise that is yet to be formally tested. Indeed, under physiologic conditions, mitochondria take up the vast majority (∼99%) of mitochondrial proteins from the cytosol via specialized import pathways.^25^ These proteins are synthesized from nuclear-encoded genes and transported across or integrated within the mitochondrial membranes^26^ and maintained via intricate PQC systems within mitochondria.^26^ Whether mitochondria take up cytosolic protein aggregates in cardiac myocytes, and whether this pathway can be therapeutically entrained for facilitating cytosolic protein homeostasis via the ‘MAGIC’ paradigm, remains unknown.

Relevant to this discussion is our recent observation where we uncovered a critical role for TRAF2, an innate immunity adaptor protein and an E3 ubiquitin ligase, as a mediator of physiologic cardiac myocyte mitophagy.^27, 28^ We discovered that inducible cardiac myocyte-specific ablation of TRAF2 markedly impairs mitophagy in the unstressed mouse heart, leading to cardiomyopathy.^27^ Importantly, it is essential to distinguish TRAF2-mediated physiologic mitophagy from stress-induced mitophagy.^28^ In the latter, a canonical pathway involving PINK1 stabilization and PARKIN recruitment to damaged mitochondria has been demonstrated to orchestrate mitophagy,^29, 30^ but PINK1 and PARKIN do not play a role in ‘basal’ or ‘physiologic’ mitophagy in unstressed adult mouse hearts.^30–32^ In the current study, we examined whether mitochondria take up aggregate-prone proteins and whether TRAF2-mediated mitophagy facilitates their removal. Our data demonstrate that TRAF2-mediated mitophagy prevents protein aggregation in the myocardium and that stimulation of mitophagy reduces protein aggregate pathology and enhances cytosolic proteostasis to restore cardiac function in proteotoxic cardiomyopathy induced by aggregate-prone proteins.

## Results

### Cardiac myocyte-specific ablation of TRAF2 provokes protein aggregation in the cytosol and in mitochondria

The discovery of mitochondrial uptake of cytosolic protein aggregates^23^ and of ubiquitinated cytosolic proteins under proteasomal inhibition stress^33^ suggests a role for mitochondria in cytosolic protein quality role, but the fate of these mitochondria-localized protein aggregates remains unknown. To examine whether mitophagy promotes removal of protein aggregates, we studied mice with inducible cardiac myocyte-specific ablation of TRAF2 (with tamoxifen treatment of *Traf2* floxed mice bearing the *Myh6*MerCreMer transgene, termed TRAF2-icKO) where we have documented impaired basal, i.e. physiologic, cardiac myocyte mitophagy,^28^ resulting in cardiomyopathy with left ventricular (LV) dilation and systolic dysfunction.^27^ We examined the presence of protein aggregates and assessed localization of DESMIN, a cytosolic scaffolding protein that is observed to localize to protein aggregates in proteotoxic cardiomyopathy.^16^ TRAF2-icKO mice demonstrate accumulation of protein aggregates in cardiac myocytes bearing p62 (Fig. 1A), an adaptor protein that binds ubiquitinated proteins and is required for formation of protein aggregates.^34^ This is accompanied by accumulation of poly-ubiquitinated proteins and p62 in the cytosolic fraction (Fig. 1B, 1D). We also observed mis-localization of DESMIN from its physiologic location at the Z-discs and the intercalated discs (see TRAF2 floxed i.e. fl/fl myocardium as control in Fig. 1A) to aggregates (see arrows in TRAF2-icKO panel in Fig. 1A). This suggests that DESMIN is ‘hijacked’ into the protein aggregates in TRAF2-icKO myocardium similar to what we have described in mice expressing an aggregate-prone R120G mutant of CRYAB.^16^ It is important to note that we observe protein aggregation in the cytosol despite intact macro-autophagy in the TRAF2-icKO mice, as we had previously demonstrated via assessment with a tandem fluorescent RFP-GFP-LC3 reporter.^27^ Furthermore, proteasome activity assayed with a fluorogenic substrate was not decreased in the TRAF2-icKO myocardium (Supplementary Figure S1) as compared with controls, demonstrating intact proteasomal function.

**Figure 1.**
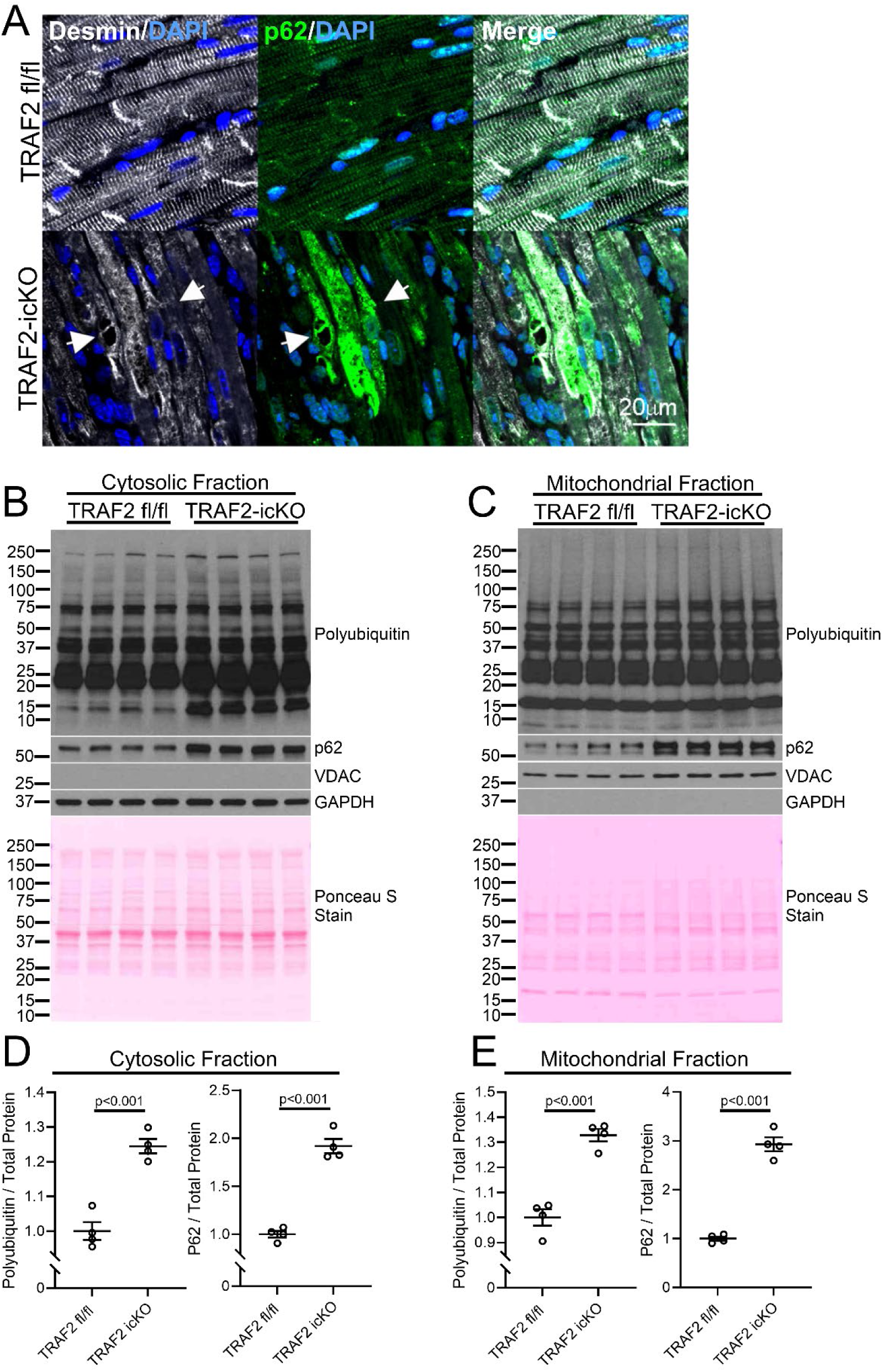
Loss of TRAF2 results in mis-localization of DESMIN in the cytosol and increased protein aggregation in the cytosol and mitochondria. **A** Representative images showing immunohistochemical staining for DESMIN (white) and p62 (green) in hearts from Traf2 floxed mice with the Myh6MerCreMer transgene (TRAF2-icKO) or Traf2 flox/flox (TRAF2 fl/fl) control mice. Adult mice were administered tamoxifen in chow for one week and sacrificed 3 weeks later. White arrows indicate DESMIN co-localized with p62-positive protein aggregates in TRAF2-icKO hearts. **B, C** Immunoblotting for protein aggregation markers p62 and polyubiquitin in the cytosolic and mitochondrial biochemical fractions from TRAF2-icKO and TRAF2 fl/fl control mouse hearts treated as in A. Immunoblotting for VDAC and GAPDH is shown to demonstrate separation of mitochondrial and cytosolic fractions, respectively. Ponceau S staining is shown to assess total protein loading. **D, E** Quantitation of the cytosolic (D) and mitochondrial (E) levels of polyubiquitin and p62 from panels B and C respectively. Polyubiquitin and p62 levels were normalized to total protein as assessed by Ponceau S Staining and expressed as a fold change relative to levels in TRAF2 fl/fl control samples. P values shown are by t-test.

Interestingly, the biochemical fractionation of TRAF2-icKO myocardium also revealed accumulation of poly-ubiquitinated proteins and p62 in the mitochondria-enriched biochemical fraction (Fig. 1C, E). Super-resolution imaging of TRAF2-icKO myocardium demonstrated co-localization of poly-ubiquitinated proteins and p62 within COXIV de-limited inner mitochondrial membrane in cardiac myocytes, which was not observed in the control mouse myocardium (Fig. 2A-C). To examine if impaired TRAF2-mediated mitophagy is sufficient to acutely induce protein aggregate accumulation within mitochondria, we isolated neonatal mouse cardiac myocytes from mice homozygous for floxed *Traf2* alleles and transduced with adenoviral Cre in cell culture to induce TRAF2 ablation. We have described that *Traf2* ablation in this setting impairs mitophagy.^27^ Loss of TRAF2 resulted in accumulation of poly-ubiquitinated proteins and p62 within mitochondria in cardiac myocytes (Fig. 2D, E). Notably, the expression of mitochondrial proteases that degrade mitochondrial protein aggregates^24^ and are known to be expressed in cardiac myocytes,^35^ namely LONP1, CLPP, HTRA2, was not significantly altered by TRAF2 ablation in the myocardium (Supplementary Fig. S2A, B), suggesting that insufficiency of mitochondrial protein degradation machinery is not the primary cause of increased mitochondrial protein aggregation in the setting of TRAF2 deletion. This suggests that mitophagy plays a unique role, separate from other protein degradation pathways, in cytosolic protein quality control in cardiac myocytes.

**Figure 2.**
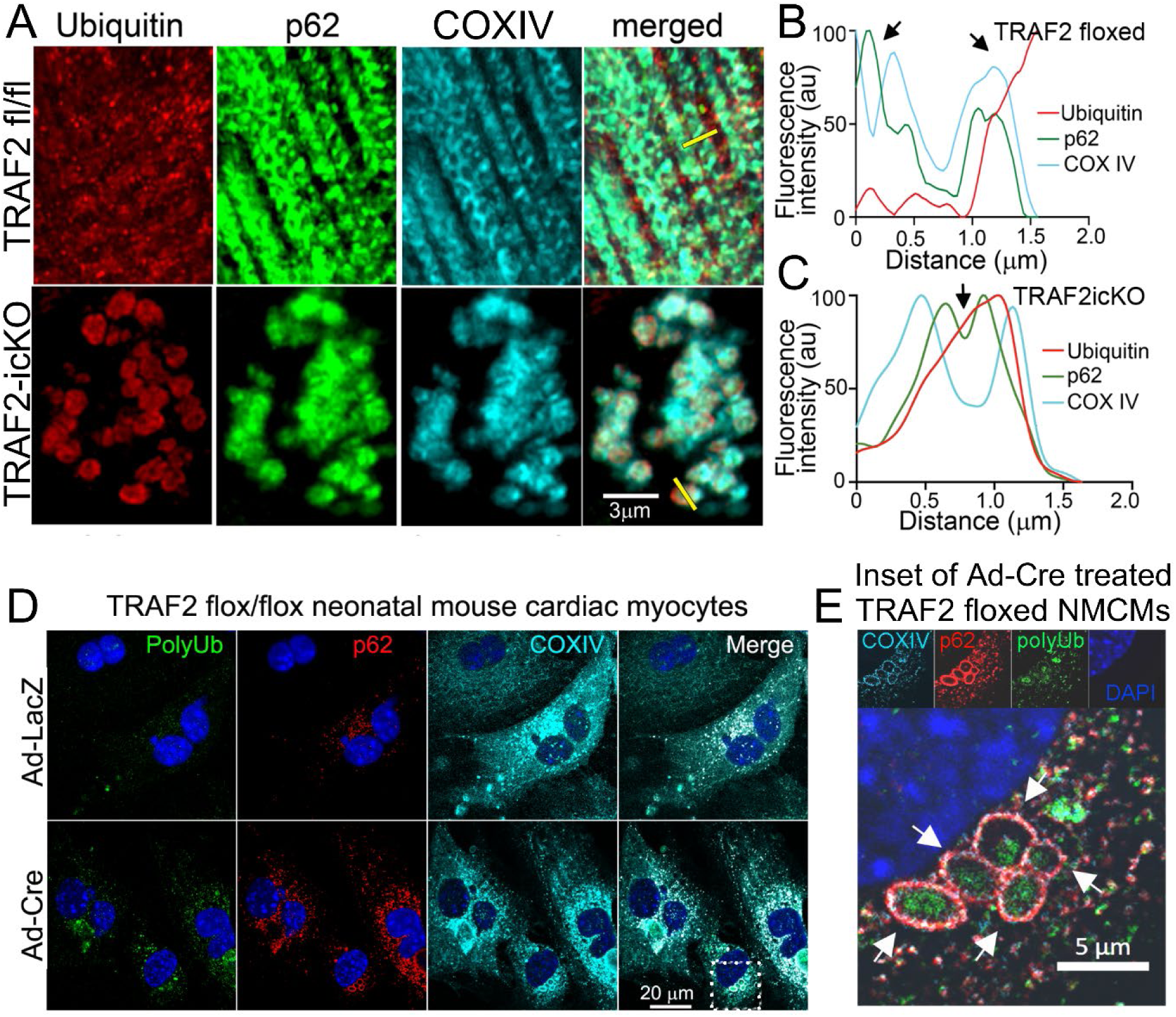
Cardiac-myocyte specific ablation of TRAF2 induces increased protein aggregation in mitochondria. **A** Representative super-resolution imaging of TRAF2-icKO and TRAF2 floxed control hearts following immunohistochemical staining for polyubiquitin (red) and p62 (green) to identify protein aggregates, and COXIV (blue) to identify the inner mitochondrial membrane. **B, C** Line scan analysis was performed along the yellow lines shown in merged images in panel A to assess the localization of p62 and polyubiquitin relative to COXIV. TRAF2 floxed hearts demonstrate p62 and polyubiquitin outside the COXIV-positive region (arrows in panel B), whereas TRAF2-icKO hearts demonstrate p62 and ubiquitin inside the region delineated with COXIV (arrows in panel C), indicating localization within the mitochondria. **D, E** Representative images showing immunohistochemical staining for polyubiquitin (green), p62 (red), and COXIV (blue) in postnatal day 1 TRAF2 floxed neonatal mouse cardiac myocytes (NMCMs) treated with adenoviral Cre (Ad-Cre) or adenoviral LacZ (Ad-LacZ) control virus, 3 days after adenoviral treatment. Arrows in magnified inset (E) from Ad-Cre treated cells (marked by dotted line in D) point to increased localization of polyubiquitinated proteins and p62 within COXIV-delineated mitochondria in NMCMs with adenoviral Cre-mediated deletion of TRAF2.

### Aggregate-prone mutants of cytosolic proteins accumulate in mitochondria

Characterization of components of aggregates induced by exogenously-expressed aggregate-prone proteins in yeast has revealed evidence for their mitochondrial uptake.^23^ Also, proteotoxic stress induced by proteasome inhibition stimulated mitochondrial uptake of poly-ubiquitinated proteins through an interaction between a previously characterized mitophagy-regulator FUNDC1 and a heat-shock chaperone protein, HSC70, on the mitochondrial outer membrane^33^. Additionally, small heat shock proteins that are essential components of the cytosolic protein quality control pathways^1^ have been detected in the inter-membranous space of the mitochondria under physiologic conditions.^36^ These proteins include CRYAB (HSPB5) and we and others have found that an arginine to glycine (R120G) mutant CRYAB protein that provokes protein aggregate pathology in cardiac myocytes^7, 16, 37^ also interacts with mitochondrial proteins^38^ and induces mitochondrial dysfunction^16, 39^. Based on these observations, we hypothesized that R120G CRYAB mutant protein will demonstrate an increased propensity to traffic to the mitochondria as compared with wild type CRYAB. We first confirmed that exogenously expressed human CRYAB R120G mutant protein forms aggregates and observed that it also co-localizes with mitochondria in HEK293 cells (Fig. 3A), consistent with prior observations in mouse cardiac myocytes.^38^ We next asked whether human cardiac myocytes exhibit similar pathology with expression of CRYAB with the R120G mutation. We generated a human induced pluripotent stem cell-derived cardiac myocyte (iPSC-CM) line homozygous for the R120G mutation in the CRYAB gene, which also harbors an mKate-α-Actinin reporter allele.^40^ iPSC-CMs homozygous for the R120G CRYAB mutation exhibited disrupted sarcomere structure relative to wild-type iPSC-CMs, as shown by expression of the mKate-α-Actinin reporter (Fig. 3B, Supplementary Figure S3). Mutant iPSC-CMs also demonstrated ubiquitin-positive structures consistent with protein aggregates (arrowheads in Fig 3B), with mis-localization of DESMIN to aggregates in myocytes matured in cell culture (Supplementary Figure S3). Isolation of cytosolic and mitochondrial fractions demonstrated that R120G mutant CRYAB preferentially localizes to the mitochondria in human iPSC-CMs (Fig 3C, D), whereas wild-type CRYAB was primarily found in the cytosolic fraction (Fig 3C, E). These findings demonstrate that the aggregate-prone R120G mutation in CRYAB leads to increased accumulation of CRYAB in the mitochondria.

**Figure 3.**
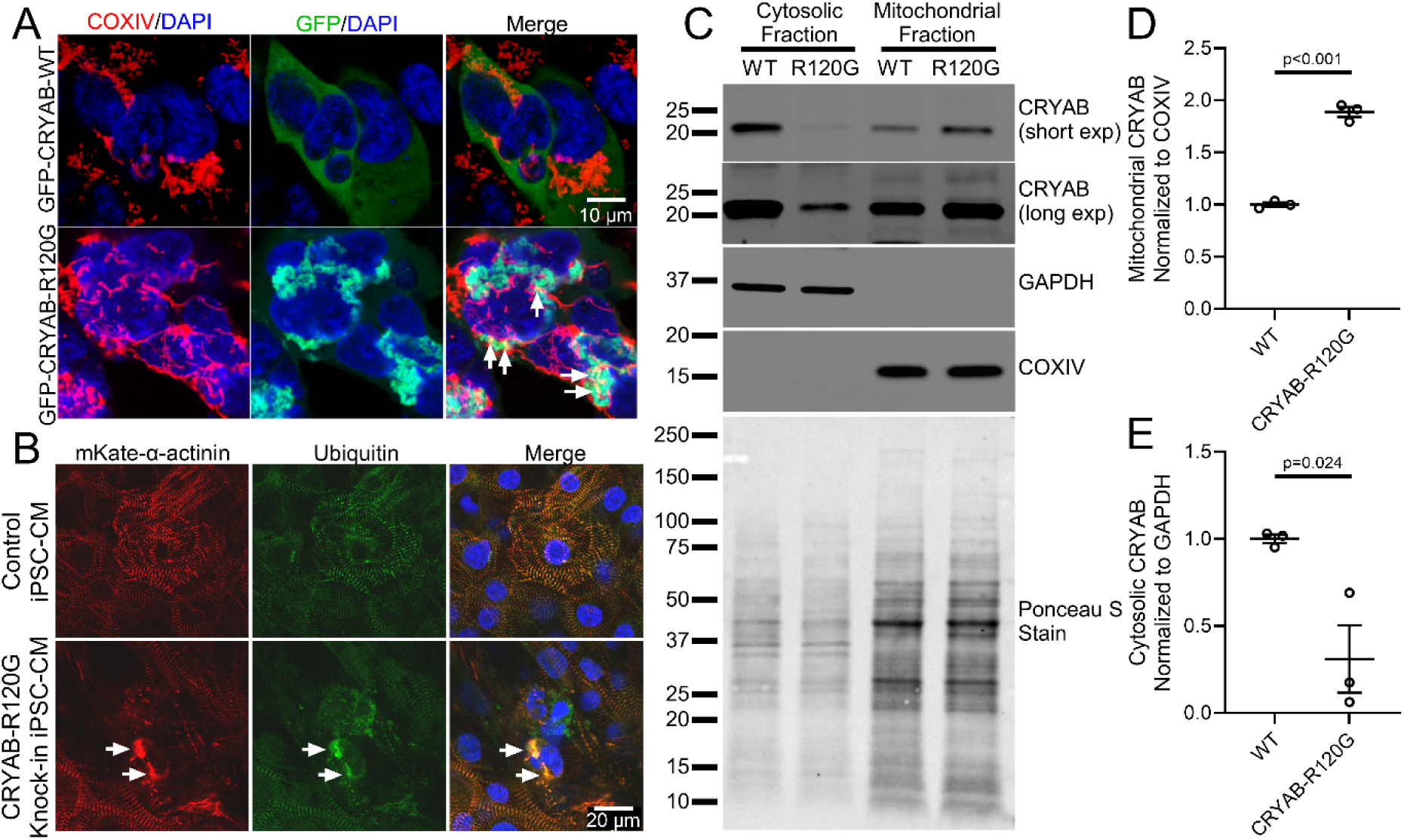
R120G mutant CRYAB protein localizes to mitochondria. **A** Representative images of immunohistochemical staining for mitochondrial marker COXIV (red) and GFP (green) in HEK293 cells transfected with GFP-tagged wild-type CRYAB (GFP-CRYAB-WT) or GFP-tagged CRYAB with the aggregation-prone R120G mutation (GFP-CRYAB-R120G). White arrows point to co-localization of COXIV with mutant GFP-CRYAB-R120G. **B** Representative images of induced pluripotent stem cell (iPSC)-derived cardiac myocytes from iPSC lines homozygous for knock-in of R120G mutation and isogenic controls expressing mKate-tagged α-actinin and stained with antibody against polyubiquitin. White arrows point to co-localization of α-actinin with polyubiquitin in aggregates. **C** Representative immunoblot demonstrating expression of CRYAB in mitochondria rich and cytosolic sub-cellular fractions from iPSC-derived cardiac myocytes homozygous for knock-in of R120G mutation (R120G) and isogenic controls (WT). GAPDH and COXIV are shown as markers for cytosol and mitochondrial fractions respectively, and Ponceau S staining is shown for protein loading control. **D, E** Quantitative evaluation of CRYAB abundance in mitochondrial (D) and cytosolic (E) fractions from experiments as shown in C, normalized as indicated and expressed as fold over control. P values shown are by t-test.

The above findings suggest that aggregate-prone mutant proteins preferentially accumulate in the mitochondrial fraction, but it remains uncertain if these proteins are imported within mitochondria or adherent on the mitochondria surface. To determine whether mitochondria take up CRYAB or its R120G mutant form, we incubated isolated mitochondria with mitochondria-depleted cellular extracts from cells expressing the GFP-tagged R120G mutant or wild type CRYAB protein, and performed a mitochondrial import assay (Fig. 4A).^41^ Subsequently, we also performed a mitochondrial protection assay with digitonin treatment to generate mitoplasts (i.e. mitochondria with intact inner membrane but stripped of the outer membrane, Fig. 4A) as previously described^33, 42^, in order to determine whether these proteins are taken up into the mitochondrial matrix rather than being associated with the outer membrane or residing in the inter-membranous space.^36^ As shown in Fig. 4B-C, while both wild-type and the R120G mutant CRYAB protein can be pulled down with isolated mitochondria, most of the R120G mutant protein (but not wild type CRYAB) is retained within digitonin-treated mitoplasts, indicating that the mutant aggregate-prone protein is taken up across the inner mitochondrial membrane into the mitochondrial matrix. To assess if this observation extends to other aggregate-prone proteins that induce cardiac proteotoxicity, we examined the P209L mutant of BAG3,^43^ which forms protein aggregates in cardiac myocytes and is implicated in causing cardiomyopathy. Indeed, as shown in Fig. 4D-E, both wild type BAG3 and the P209L mutant can be pulled down with isolated mitochondria, but only the P209L mutant (but not the wild type BAG3 protein) is retained after digitonin treatment indicating that it was taken up across the inner mitochondrial membrane into the mitochondrial matrix. Prior work has demonstrated that preservation of mitochondrial membrane potential is necessary for import of mitochondrial proteins^23, 41^. To test whether intact mitochondrial membrane potential was required for the import of CRYAB-R120G, we pre-treated mitochondria with the mitochondrial uncoupler CCCP to disrupt membrane potential prior to performing import assays and generating mitoplasts via digitonin treatment. Pre-treatment of mitochondria with CCCP significantly reduced import of CRYAB-R120G (Fig 4F, G), indicating that intact mitochondrial membrane potential is required for import of CRYAB-R120G. siRNA-mediated knockdown of HSC70 (Supplementary Figure S4A, B) resulted in a reduction in the mitochondrial import of CRYAB-R120G (Fig 4H, I). Taken together, these experiments demonstrate that aggregate-prone R120G mutant CRYAB is directly imported into the mitochondria via an HSC70-dependent process that requires intact mitochondrial membrane potential.

**Figure 4.**
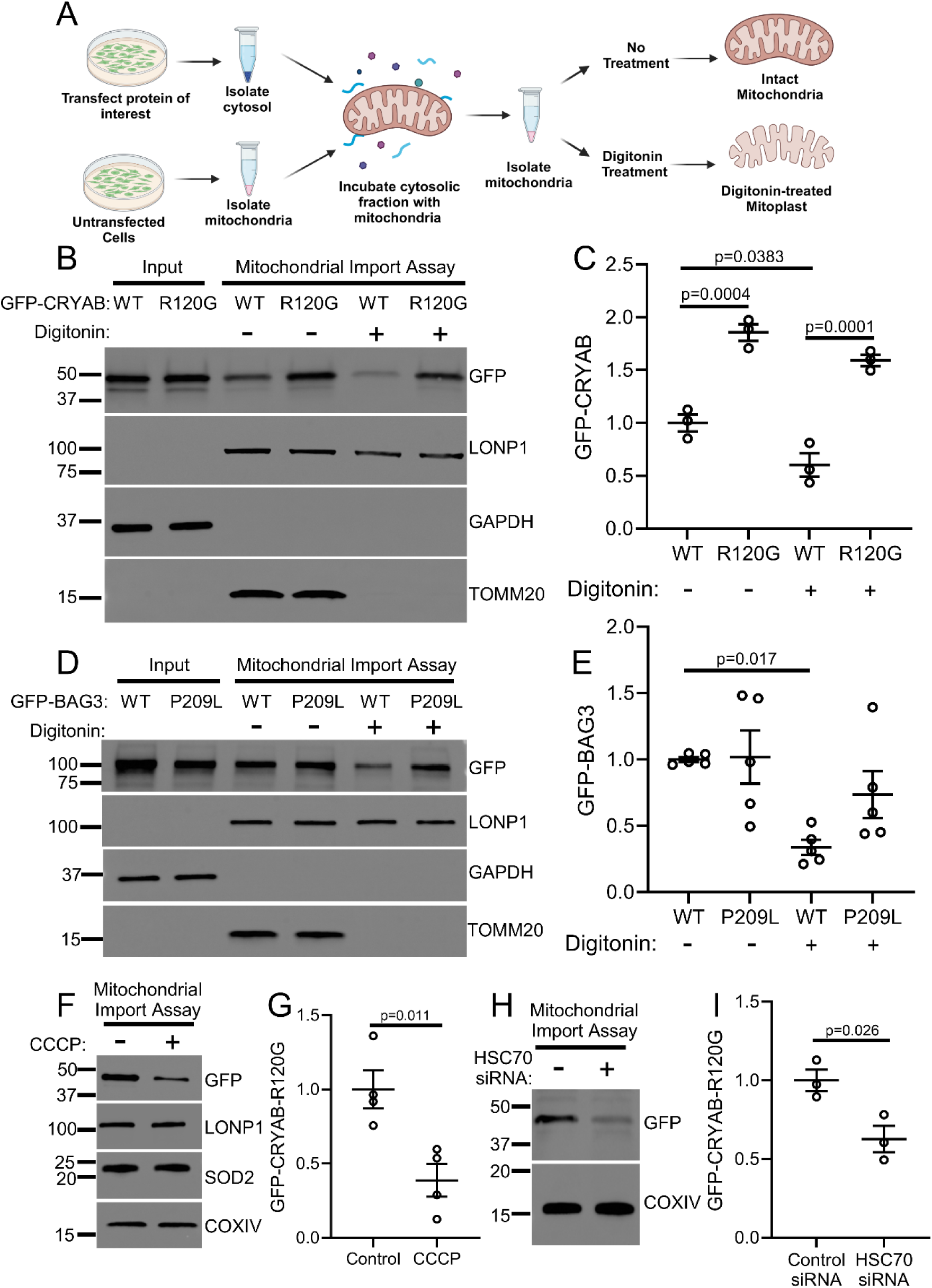
Aggregate-prone mutant proteins accumulate in the mitochondria. **A** Schematic depicting experiments for mitochondrial uptake of exogenously expressed proteins in HEK293 cells followed by mitochondrial protection assay. See methods for details. **B, C** GFP-CRYAB-WT or GFP-CRYAB-R120G were expressed in HEK293 cells and mitochondria-free cytosolic fractions were generated (as shown in input lanes). Cytosolic fractions were then incubated with isolated mitochondria from untransfected cells. Half of the mitochondrial pellet was treated with digitonin, shown as ‘+’ (and the other half with diluent shown as ‘-’) to disrupt the outer mitochondrial membrane and assessed by immunoblotting to determine the import of GFP-CRYAB into the mitochondria. GFP-CRYAB-R120G demonstrated increased association with mitochondria in comparison to GFP-CRYAB-WT, with less depletion by digitonin treatment (quantitated in C and expressed as fold change over control), indicating increased mitochondrial import of CRYAB-R120G and its retention within mitoplasts as compared with CRYAB-WT. P values shown are by one-way ANOVA followed by Tukey’s post-hoc testing. **D, E** GFP-BAG3-WT and mutant GFP-BAG3-P209L were expressed in HEK293 cells and evaluated in mitochondrial import and digitonin protection assays in analogous manner to CRYAB in panels b and c. BAG3-WT but not the mutant GFP-BAG3-P209L protein exhibited depletion upon digitonin treatment (quantitated in E and expressed as fold change over control), indicating import and retention of BAG3-P209L (but not BAG3-WT) within mitoplasts. P value shown is by one-way ANOVA followed by Tukey’s post-hoc testing. **F, G** Representative immunoblot (F) depicting expression of GFP-tagged R120G CRYAB with quantitation of abundance (G, expressed as fold change over control) in mitoplasts generated after mitochondria import assay into mitochondria pretreated with 4mM CCCP for 5min. P value shown is by t-test. **H, I** Representative immunoblot (H) depicting expression of GFP-tagged R120G CRYAB with quantitation of abundance (I, expressed as fold change over control) in mitoplasts generated after mitochondria import assay into mitochondria from cells treated with siRNA targeting HSC70 or control. P value shown is by t-test.

### Aggregate-prone R120G CRYAB mutant protein localizes to the mitochondria and induces mitophagy in the mouse myocardium

To evaluate the role of mitophagy in cardiac proteotoxicity observed with expression of aggregate-prone proteins, we studied transgenic mice (TG) with *Myh6*-promoter driven cardiac myocyte-specific expression of the human R120G CRYAB mutant protein that provokes protein aggregation, cardiac dysfunction and early mortality.^16, 37^ Intriguingly, *Myh6*-R120G CRYAB transgenic myocardium from 20-24 week-old mice demonstrates upregulation of TRAF2 (Fig 5A, B) at a stage when extensive protein aggregate pathology is evident.^37^ Next, we performed biochemical fractionation of myocardial tissue from these mice. In addition to accumulation of poly-ubiquitinated proteins and p62 (both markers of aggregates) in the cytosolic fraction (Fig. 5E), we found that a substantial fraction of the overexpressed CRYAB protein localizes to the mitochondrial fraction (Fig. 5E) accompanied by marked accumulation of p62 and poly-ubiquitinated proteins in the mitochondria as well (Fig. 5E, G, H). This is associated with increased TRAF2 localization to the mitochondrial fraction (Fig. 5E, F) and a reduction in abundance of mitochondrial proteins (VDAC and COXIV, Fig. 5A, C, D) in crude heart extracts, suggesting that cardiac myocyte mitophagy may be upregulated in these mice at this age. In agreement with these findings, prior work has shown that the content of mitochondrial enzymes such as citrate synthase and respiratory chain complexes is decreased in transgenic mice expressing the R120G CRYAB mutant protein compared with wild type controls, even prior to development of cardiomyopathy.^39^ Accordingly, we examined mitophagy in the CRYAB R120G transgenic myocardium in mice expressing the mito-QC allele, a reporter for mitophagy.^44^ We detected a significant increase in red-only fluorescent puncta within cardiac myocytes, indicating the increased presence of mitochondria in acidified organelles (wherein GFP fluorescent is quenched), demonstrating increased mitophagic sequestration of mitochondria in the setting of proteotoxic stress (Fig. 5I, J). In addition to increased expression of TRAF2, we also noted increased expression and mitochondrial localization of the canonical mitophagy mediator PARKIN (Supplementary Figure S5A-D), albeit to a lesser extent than what we observed for TRAF2 (Fig. 5E, F). Taken together, these results suggest that proteotoxic stress in cardiac myocytes induces mitophagy as a compensatory response, possibly mediated via increased expression and mitochondrial localization of TRAF2 and PARKIN.

**Figure 5.**
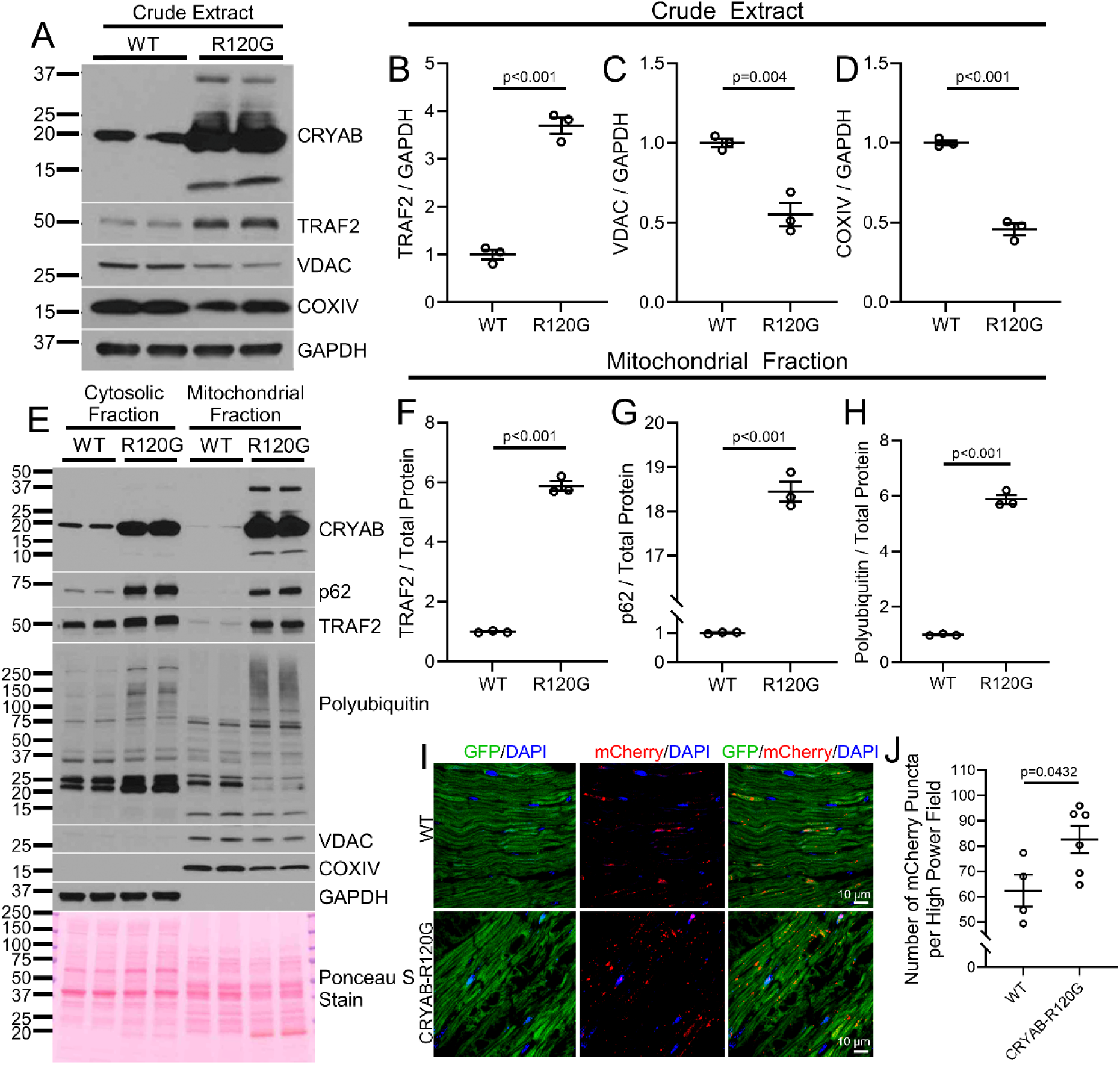
Transgenic CRYAB-R120G mice exhibit mitochondrial CRYAB localization with upregulation of TRAF2 and induction of mitophagy. **A-D** Representative immunoblot (A) demonstrating expression of CRYAB, TRAF2 and mitochondrial proteins, namely VDAC and COXIV, in crude whole heart extracts from 20-24 week-old CRYAB-R120G mice (R120G) or littermate controls (WT). R120G hearts exhibited increased TRAF2, reduced VDAC, and reduced COXIV (quantitated in B, C, and D, respectively) versus wild-type. For quantitation, samples were normalized to GAPDH to control for loading. P values shown are by t-test. **E-H** Representative immunoblot (E) on cytosolic and mitochondrial fractions from heart tissue from 20-24 week-old CRYAB-R120G mice (R120G) or littermate controls (WT), demonstrating expression of CRYAB, TRAF2, p62, and polyubiquitinated proteins. GADPH was used to demonstrate separation of the cytosolic fraction, and VDAC and COXIV were used as markers of the mitochondrial fraction. Ponceau S staining is shown as loading control. Mitochondrial fractions from R120G hearts demonstrated increased levels of the protein aggregation markers polyubiquitin and p62, as well as TRAF2 (quantitated in F, G, and H). For quantitation, samples were normalized to total protein levels measured by Ponceau S staining. P values shown are by t-test. **I, J** Mitophagy was assessed using the Mito-QC reporter allele in 20-week-old CRYAB-R120G mice (R120G) and littermate controls (WT). Mito-QC reporter expression results in dual tagging of mitochondria with GFP and mCherry signals, with GFP signal predominating in the cytoplasmic neutral pH environment and mCherry signal predominating in lysosomal acidic pH environment. Frozen sections were analyzed for GFP and mCherry signal. CRYAB-R120G hearts exhibited increased mCherry-only puncta, indicating increased mitochondrial localization to the lysosome and increased mitophagy. Panel J shows quantitation of mCherry puncta per high power field. P values shown is by t-test.

### TRAF2 deficiency worsens cardiac function in the setting of R120G CRYAB-induced proteotoxicity

The combination of dramatically increased myocardial TRAF2 expression, increased mitochondrial localization of TRAF2, and increased mitophagy seen in the hearts of R120G CRYAB transgenic mice (Fig. 5) suggests a protective role for TRAF2-induced mitophagy in response to proteotoxic stress. To test this hypothesis, we generated *Myh6*-R120G CRYAB transgenic mice with inducible ablation of one *Traf2* floxed allele with tamoxifen treatment in mice also carrying the Mer-Cre-Mer transgene expressed via the *Myh6* promoter (labeled as R120G TRAF2-icHET). We chose to ablate a single *Traf2* allele to avoid the confounding effects of cardiomyopathy with ablation of both alleles as we and others have described.^27, 45^ Loss of one *Traf2* allele is well tolerated in the mouse myocardium.^45^ Inducible adult-onset ablation of one *Traf2* allele in R120G TG mice from 8 weeks of age (Fig. 6A) resulted in worsening of left ventricular systolic function (Fig. 6B, C) at 20 weeks of age without affecting left ventricular end-diastolic dimension (Supplementary Figure S6A). R120G TRAF2-icHET mice demonstrate markedly accelerated mortality as compared with R120G CRYAB transgenic mice with intact *Traf2* alleles (Fig. 6D); while ablation of one *Traf2* allele did not affect survival over the same period in wild type controls (Supplementary Figure S6B). While myocardial histology and fibrosis were not affected at 20 weeks of age (Fig. 6E), R120G TRAF2-icHET mice did develop worsening mitochondrial morphology with worsening cristal effacement on transmission electron microscopy analysis of cardiac myocytes (Fig. 6F, see white arrows). Interestingly, this was accompanied by appearance of aggregates within mitochondria in R120G TRAF2-icHET myocardium (Fig. 6F, see black arrowheads). Inducible adult-onset ablation of one *Traf2* allele resulted in 32% reduction in cytosolic TRAF2 protein abundance (Fig. 7A, B). This was accompanied by increased accumulation of polyubiquitinated proteins in the mitochondrial fraction of R120G TRAF2-icHET mouse hearts (Fig. 7A, C). These findings indicate that cardiac myocyte upregulation of TRAF2 is protective in the setting of cardiomyopathy induced by proteotoxic stress from aggregate-prone R120G CRYAB expression, presumably through increased mitophagy. As we had observed that R120G CRYAB transgenic mice also exhibit increased PARKIN expression (Supplementary Figure S5), we next examined whether loss of PARKIN would also worsen the cardiomyopathy in these mice, by generating R120G CRYAB mice that were also homozygous for the *Park2* null allele and therefore deficient in PARKIN (R120G-CRYAB-PARKIN KO). Surprisingly, global loss of PARKIN did not affect left ventricular systolic function or end-diastolic dimension in *Myh6*-R120G CRYAB transgenic mice (Supplementary Figure S7A, B). These results indicate that PARKIN does not play a protective role in the cardiomyopathy seen in R120G CRYAB transgenic mice and suggest that the protective effects of mitophagy in response to proteotoxic stress are PARKIN-independent, as has been observed previously in both stressed and unstressed mouse myocardium.^30–32, 46–48^

**Figure 6.**
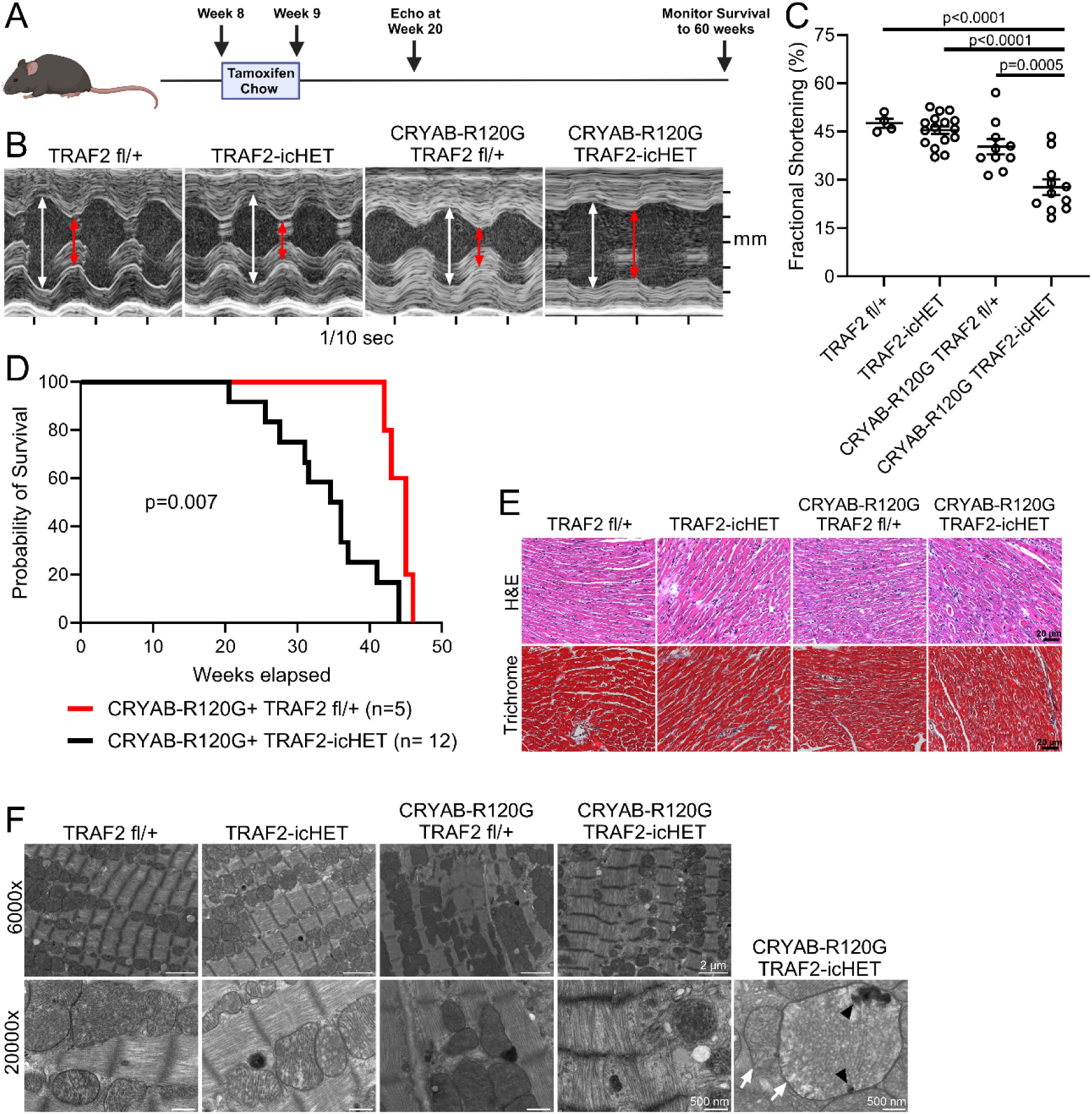
Reduction in cardiac myocyte TRAF2 exacerbates CRYAB-R120G-induced cardiomyopathy. **A** Experimental approach to deplete TRAF2 in cardiac myocytes. One Traf2 floxed allele and the cardiac myocyte-specific MerCreMer tamoxifen-inducible Cre allele were introduced into CRYAB-R120G transgenic line to generate CRYAB-R120G mice with cardiac myocyte-specific inducible deletion of one Traf2 allele (termed R120G TRAF2-icHET). These mice were compared to R120G TRAF2 fl/+ mice (without the MerCreMer transgene) and to TRAF2 fl/+ and TRAF2-icHET control mice without CRYAB-R120G. Mice were treated with tamoxifen chow for one week starting at 8 weeks of age. Echocardiography was done at 20 weeks of age, and survival was monitored to 60 weeks. **B** Representative M-mode echocardiograms for 20-week-old mice in indicated groups. White arrows indicate left ventricular internal diastolic diameter, red arrows indicate left ventricular internal systolic diameter. **C** Assessment of left ventricular systolic function by fractional shortening in 20-week-old mice. Fractional shortening was reduced in R120G TRAF2-icHET mice versus R120G TRAF2 fl/+ controls. P values shown are by one-way ANOVA followed by Tukey’s test for multiple comparison testing between groups. **D** Kaplan-Meier survival analysis of R120G TRAF2-icHET mice, which exhibited earlier mortality versus CRYAB-R120G TRAF2 fl/+ mice. P value shown is by Mantel-Cox log-rank testing. **E** Representative hematoxylin and eosin, and Masson’s trichrome-stained images of hearts from 20-week-old mice from indicated groups modeled as in A. **F** Representative transmission electron microscopy of heart tissue from 20-week-old mice from indicated groups. CRYAB-R120G TRAF2-icHET hearts exhibit disrupted mitochondrial morphology with cristal rarefaction (white arrows) and mitochondrial protein aggregates (black arrowheads).

**Figure 7.**
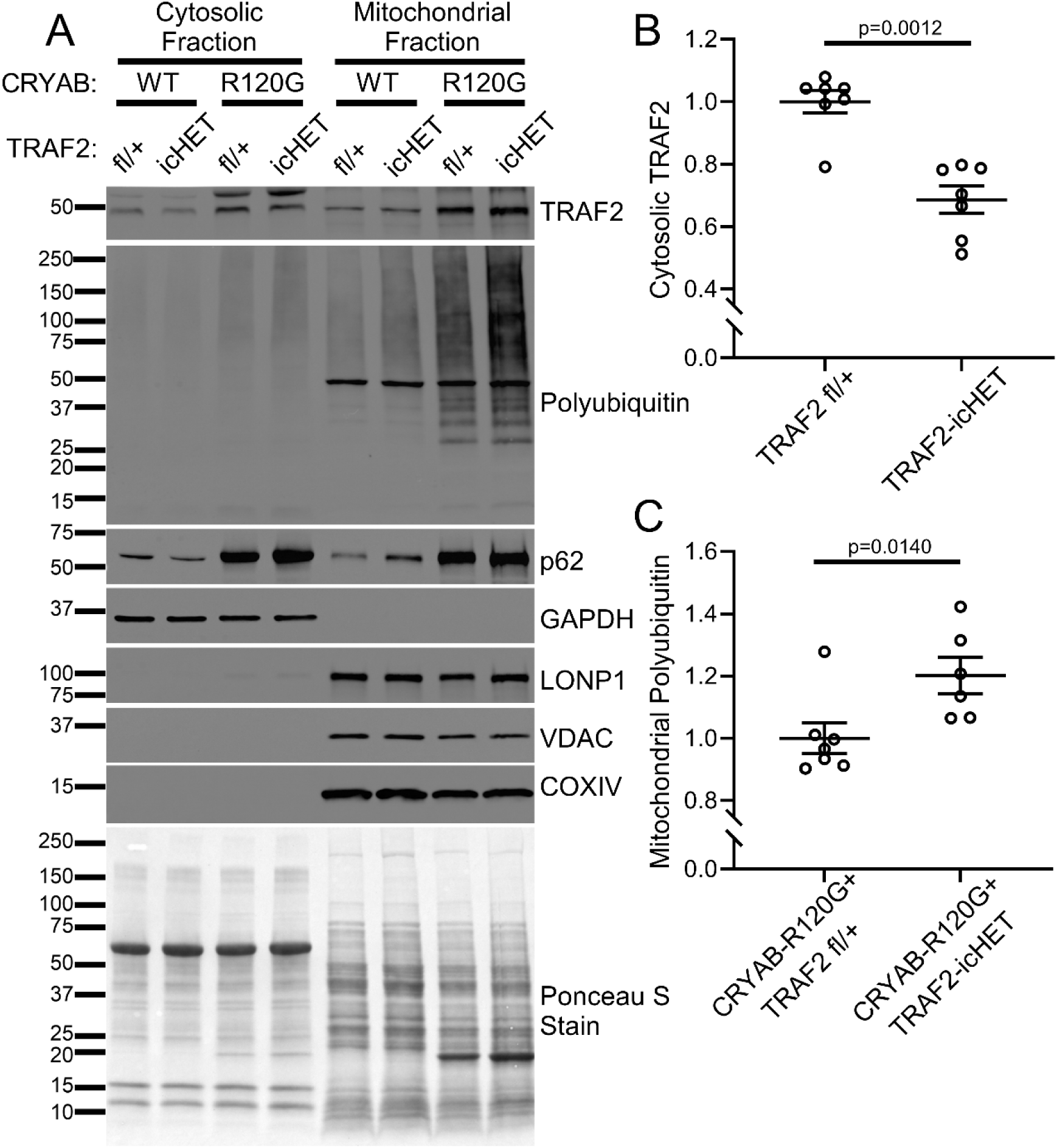
Reduction in cardiac myocyte TRAF2 increased mitochondrial protein aggregates in R120G transgenic myocardium. **A** Representative immunoblot depicting expression of TRAF2, polyubiquitinated proteins and p62 in cytosolic and mitochondrial fractions from 20-week-old CRYAB-R120G mice with cardiac myocyte-specific inducible deletion of one Traf2 allele at 8 weeks of age (termed R120G TRAF2-icHET), and similarly treated R120G TRAF2 fl/+ mice, TRAF2-icHET and TRAF2 fl/+ as control as modeled in Figure 6A. GADPH was used to demonstrate separation of the cytosolic fraction, and LONP1, VDAC and COXIV were used as markers of the mitochondrial fraction. **B** Quantitation of TRAF2 levels in the cytosolic fractions from TRAF2-icHET and TRAF2 fl/+ mice treated as in A. P value shown is by Mann-Whitney test. **C** Quantitation of polyubiquitinated proteins in the mitochondrial fractions from R120G TRAF2-icHET and R120G TRAF2 fl/+ mice treated as in A. P value shown is by Mann-Whitney test.

### Targeted TRAF2 overexpression rescues mortality and cardiac function in the setting of R120G CRYAB-induced proteotoxicity

To test the hypothesis that further augmentation of mitophagy ameliorates proteotoxicity induced by the aggregate-prone R120G mutant CRYAB protein, we performed AAV9-mediated transduction of TRAF2 protein (driven by the cardiac troponin T promoter to achieve cardiac myocyte specific targeting^27^) or GFP as control in adult 8-week-old *Myh6*-R120G CRYAB transgenic mice (Fig. 8A). Our prior published studies have demonstrated that this strategy is sufficient to induce mitophagy in the mouse myocardium.^27^ TRAF2 transduction in cardiac myocytes significantly improved left ventricular systolic function (Fig. 8B, C) in AAV9-cTnT-TRAF2 transduced R120G transgenic mice as compared with AAV9-cTnT-GFP transduced R120G transgenic mice (as control) at 20 weeks of age (without a change in left ventricular end-diastolic dimension (Supplementary Fig. S8A)). Importantly, AAV9-mediated transduction of TRAF2 significantly delayed mortality in the *Myh6*-R120G CRYAB transgenic mice (Fig. 8D) where markedly accelerated mortality is observed at ∼40 weeks of age,^16, 37^ without affecting mortality in wild type mice as compared with AAV9-GFP group (Supplementary Fig. S8B). This was associated with improvement in mitochondrial morphology on transmission electron microscopy analysis of cardiac myocytes (Fig. 8F) without a discernible change in myocardial histology or fibrosis (Fig. 8E), suggesting that TRAF2-mediated benefits were transduced via its effects on the mitochondria. Indeed, AAV9-mediated transduction of TRAF2 improved mitochondrial function in the *Myh6*-R120G CRYAB transgenic mice, as assessed with oxygen consumption via respiratory chain complex I and II, and maximal respiration which are observed to be reduced in *Myh6*-R120G CRYAB transgenic mice when compared with controls (Fig. 8G, H; Supplementary Figure S9A-C), as described previously.^39^

**Figure 8.**
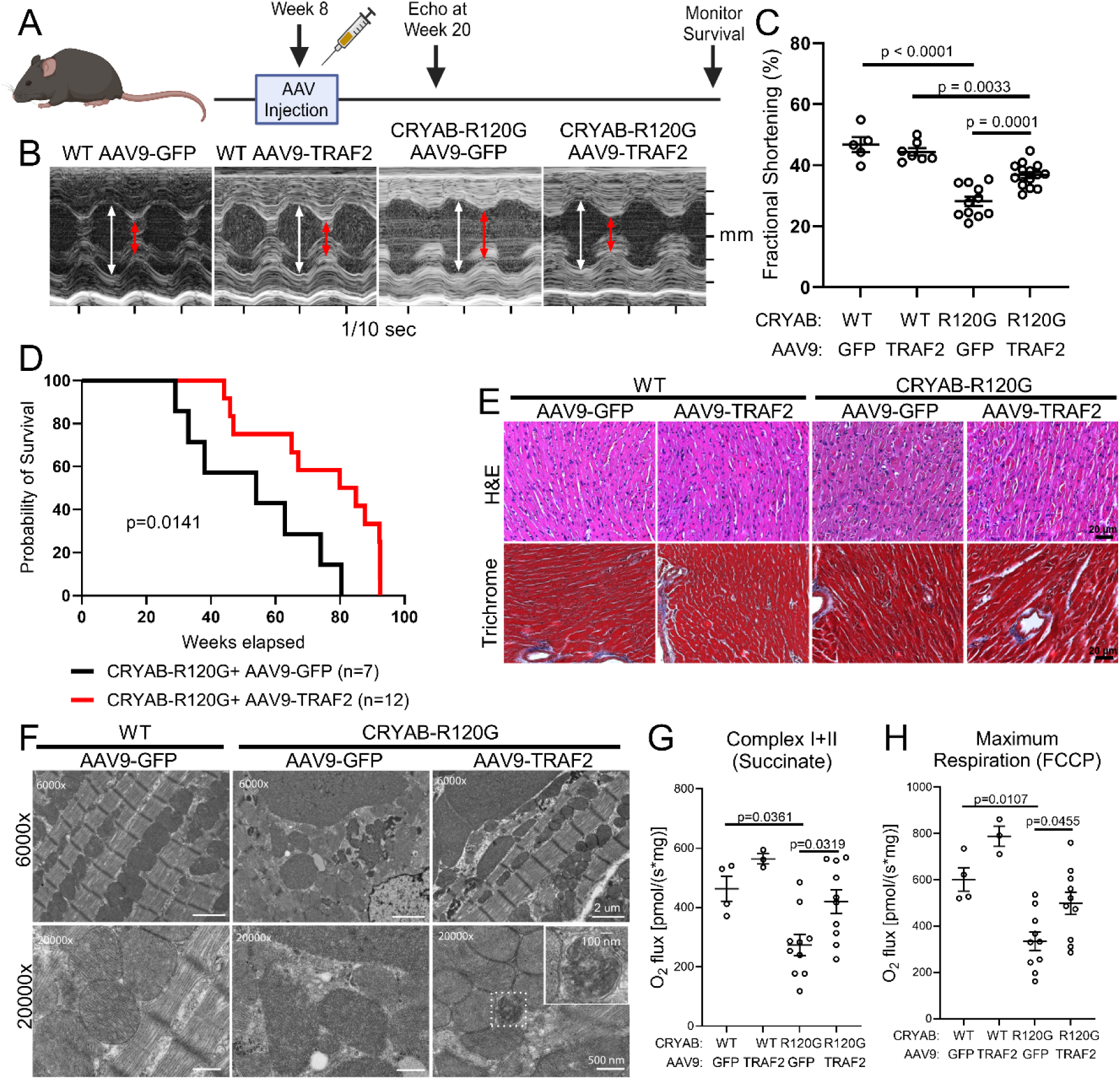
TRAF2 overexpression rescues CRYAB-R120G-induced cardiomyopathy. **A** Experimental approach to overexpress TRAF2 in cardiac myocytes. Wild-type (WT) and transgenic CRYAB-R120G mice were injected with AAV9 expressing TRAF2 (AAV9-TRAF2) or control AAV9 expressing GFP (AAV9-GFP). Cardiac troponin T promoter was employed in both viruses to specifically target cardiac myocytes. AAVs were injected via tail-vein at a dose of 7.0 X 10^11^ viral genomes/mouse at 8 weeks of age. Echocardiography was performed at 20 weeks, and survival was monitored to 100 weeks. **B** Representative M-mode echocardiograms for 20-week-old mice in indicated groups modeled as in A. White arrows indicate left ventricular internal diastolic diameter; red arrows indicate left ventricular internal systolic diameter. **C** Assessment of systolic function by fractional shortening in 20-week-old mice. Left ventricular fractional shortening is improved in CRYAB-R120G mice treated with AAV9-TRAF2 versus AAV9-GFP. P values are by one-way ANOVA followed by Tukey’s test for multiple comparison testing between groups. **D** Survival of WT and CRYAB-R120G mice injected with AAV9-GFP and AAV9-TRAF2 viruses. CRYAB-R120G AAV9-TRAF2 mice exhibited improved survival versus CRYAB-R120G AAV9-GFP mice. P value shown is by Mantel-Cox log-rank testing. **E** Representative hematoxylin and eosin and Masson’s trichrome-stained images of hearts from 20-week-old mice from indicated groups modeled as in A. **F** Transmission electron microscopy of heart tissue from 20-week-old mice from indicated groups. AAV9-TRAF2 treatment improved mitochondrial morphology in CRYAB-R120G mice relative to AAV9-GFP treatment in these animals. Increased prevalence of mitochondria sequestered within double membranes, suggesting mitochondria undergoing mitophagy, was also seen in CRYAB-R120G AAV9-TRAF2 hearts (see inset). **G, H** High-resolution respirometry was performed to measure oxygen consumption [volume-specific oxygen flux (JO2)] in isolated, permeabilized cardiac fiber bundles from mice modeled as in A in the presence of succinate (to assess complex I+II in g) and FCCP (to assess maximal respiration in h). P values are by one-way ANOVA followed by Tukey’s post-hoc test for multiple comparison testing.

To examine if TRAF2 transduction results in improvement of myocardial protein aggregate pathology, we performed subcellular fractionation of AAV9-cTnT-TRAF2 (or AAV9-cTnT-GFP) transduced R120G CRYAB TG myocardium at 20 weeks of age. AAV9-mediated TRAF2 transduction resulted in increased abundance of cytosolic and mitochondrial TRAF2 (Fig. 9A, B, C) with reduction in the abundance of poly-ubiquitinated proteins and p62 (Fig. 9A, D, E) in the mitochondrial fraction from *Myh6*-R120G CRYAB transgenic mouse hearts as compared with AAV9-cTnT-GFP-treated *Myh6*-R120G CRYAB transgenic as control. This indicates a reduction in mitochondrial protein aggregates, presumably through TRAF2-induced acceleration of mitophagy. Furthermore, AAV9-cTnT-TRAF2 treatment resulted in DESMIN re-localization to normal physiologic location on Z-discs and intercalated discs, with reduction in myocardial protein aggregates as compared with control AAV9 treatment in the R120G TG myocardium (Fig. 9F, G). Taken together, these findings suggest that cardiac myocyte mitophagy plays a protective role in removal of toxic aggregate-prone R120G mutant of CRYAB resulting in restoration of DESMIN localization in the cytosol and the preservation of cardiac function.

**Figure 9.**
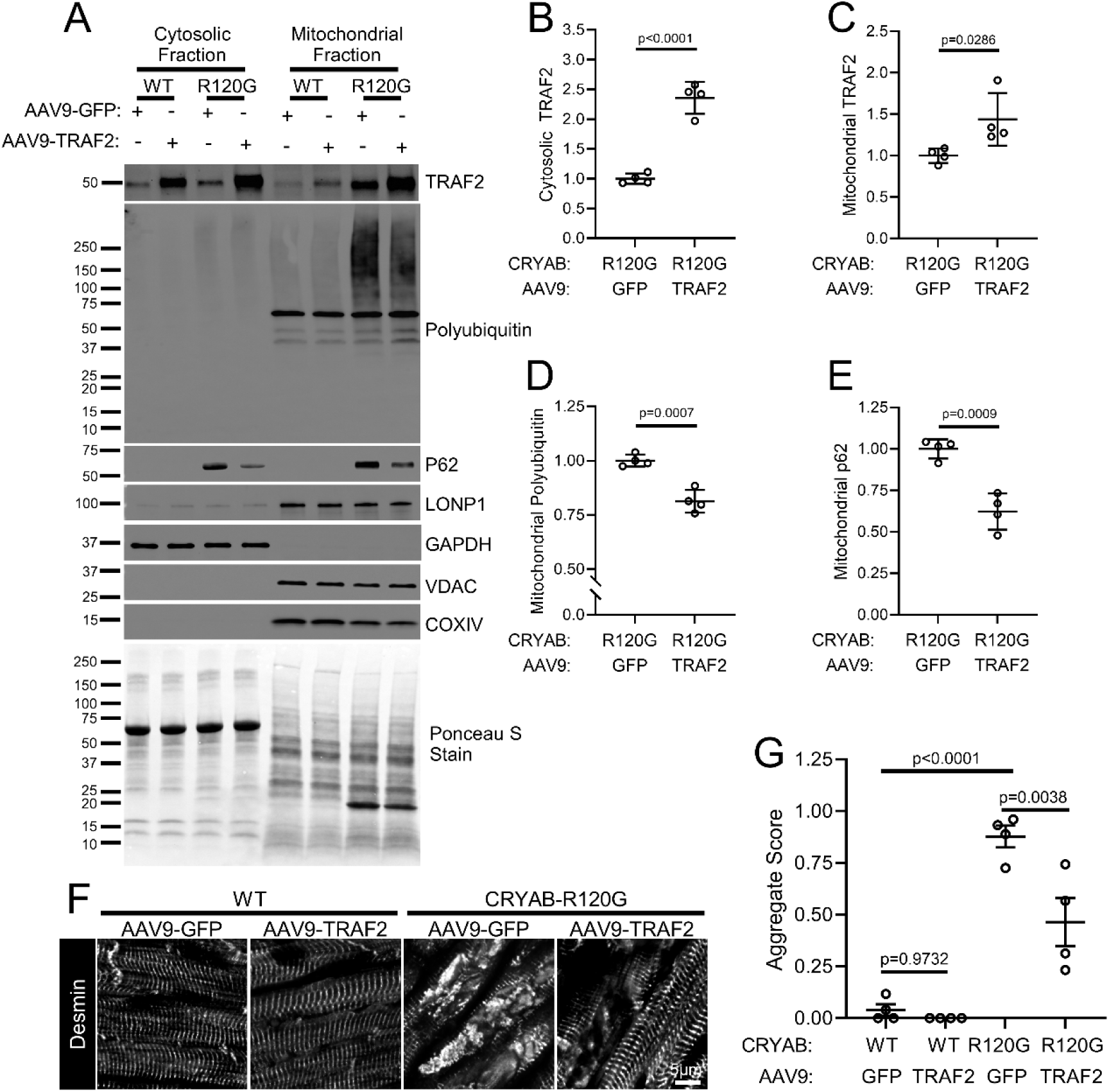
TRAF2 overexpression improves mitochondrial protein aggregation and normalizes DESMIN localization in CRYAB-R120G transgenic hearts. **A-E** Representative immunoblots (A) of cytosolic and mitochondrial fractions from 20-week-old wild-type (WT) AAV9-GFP, WT AAV9-TRAF2, CRYAB-R120G AAV9-GFP, and CRYAB-R120G AAV9-TRAF2 hearts. Mice were injected with AAV virus at 8 weeks, as previously described in Figure 8A. Quantitation of cytosolic TRAF2, mitochondrial TRAF2, mitochondrial polyubiquitin, and mitochondrial p62 levels are shown in B, C, D, and E respectively, with normalization to total protein level as assessed by Ponceau S staining. Values shown as fold change over control. P values are by t-test for B, D and E; and by Mann-Whitney test for panel C. **F** Representative images depicting immunohistochemical analysis to assess DESMIN localization in cardiomyocytes. CRYAB-R120G AAV9-GFP hearts exhibited disruption of normal striated DESMIN pattern seen in WT AAV-GFP and WT AAV9-TRAF2 mice, with localization of DESMIN to aggregates. DESMIN localization (pseudo colored in white) was partially rescued with reduced aggregates in CRYAB-R120G AAV9-TRAF2 mice. **G** Quantitative assessment of protein aggregates seen by histology in mice treated as in A at 20 weeks of age. P values are by one-way ANOVA followed by Tukey’s post-hoc test.

## Discussion

Mitochondria sustain cardiac function by generating the energy required for contraction.^49^ Also, mitochondrial integrity controls critical life and death decisions, and regulates inflammation in cardiac physiology and pathology.^49, 50^ Our findings provide evidence for another critical function for mitochondria in cardiac myocytes, namely to facilitate protein quality control in the cytosol to maintain sarcomere homeostasis. The following lines of evidence support this claim. First, impairment of physiologic cardiac myocyte mitophagy with ablation of TRAF2 induces accumulation of protein aggregates in the cytosol (along with protein aggregates in the mitochondria) and mis-localizes DESMIN to protein aggregates. Second, aggregate-prone proteins that induce cardiomyopathy and heart failure in humans, namely the R120G mutant of CRYAB and the P209L mutant of BAG3, are actively taken up by the mitochondria and co-localize with mitochondrial matrix proteins. Third, a mouse model of R120G CRYAB protein aggregate-induced cardiomyopathy demonstrates presence of mutant CRYAB protein in the mitochondria with upregulation of mitophagy prior to development of left ventricular systolic dysfunction. Fourth, attenuation of mitophagy via reduction of TRAF2 markedly accelerates development of cardiomyopathy with increased mortality in these mice. Lastly, mitophagy can be stimulated via TRAF2 overexpression to attenuate functional decompensation, delay mortality, attenuate mitochondrial protein aggregates and restore normal DESMIN localization in mice with R120G-induced cardiac proteotoxicity. Taken together, these findings lead us to propose a novel paradigm in PQC mechanisms that control sarcomere integrity. Cardiac mitochondria occupy prime real estate next to the sarcomeres and are well-positioned to regulate sarcomere structure through facilitating protein quality control. They also provide readily accessible capacity to sequester mis-folded proteins and remove them through mitophagy, a function that can be scaled up under stress to sustain sarcomere protein quality and function.

Our findings point to a critical role for mitochondria as ‘early responders’ to cytosolic proteotoxic stress in cardiac myocytes. Under these conditions, mitochondria upregulate sequestration of aggregate-prone proteins and facilitate their removal. Future studies will focus on understanding the mechanisms for uptake of protein aggregates in cardiac mitochondria, vis-à-vis the mechanisms of physiologic protein import into mitochondria.^25^ Our observations indicate that healthy mitochondria can take up aggregate-prone proteins, namely the R120G mutant of CRYAB and the P209L mutant of BAG3. Questions remain regarding whether the uptake of aggregate-prone proteins is an energy-requiring process, and whether the mechanisms of their uptake parallel those for import of normal mitochondrial proteins in physiology.^25, 51^ Our data indicate that stimulation of mitophagy facilitates the removal of a fraction of aggregates in the R120G CRYAB transgenic mice. Identifying the pathways that promote mitochondrial uptake of misfolded cytosolic proteins offers a therapeutic opportunity to increase the contribution of mitochondrial PQC machinery and mitophagy in attenuating cytosolic protein aggregate pathology.

Our data suggest that imported aggregate-prone proteins overwhelm the mitochondrial protein quality control machinery^26^ and form aggregates within the mitochondria to induce mitochondrial damage and dysfunction. Prior observations indicate that R120G mutant of CRYAB localizes to the mitochondria by interacting with an inner membrane protein VDAC^38^ along with DESMIN. Studies from our lab^16^ and others^39^ show that mitochondria are damaged and dysfunctional in cardiac myocytes expressing the R120G CRYAB mutant protein in cell culture,^16^ and in transgenic mice expressing the R120G mutant form of human or murine protein in mouse cardiac myocytes.^16, 38, 39^ These mitochondrial abnormalities, which include alterations in their organization and ultrastructure,^16, 38, 39^ reduction in respiratory chain complexes and reduced oxygen consumption,^38, 39^ and loss of mitochondrial inner membrane potential and activation of cell death signaling pathways^16, 38^ accompany the appearance of protein aggregate pathology and precede the development of cardiomyopathy and heart failure.^16, 38, 39^ Also, mitochondrial levels of Drp1, as well as its Ser616 phosphorylated from (i.e. the active form) were increased and levels of Opa1 were reduced in transgenic mice expressing R120G CRYAB mutant protein in cardiac myocytes;^39^ indicating a shift in balance towards mitochondrial fission that precedes sequestration of mitochondria by mitophagy.

Our data also suggest that mitophagy is stimulated via upregulation of TRAF2 protein and translocation of TRAF2 protein to the mitochondria in the setting of protein aggregate pathology, and that induction of mitophagy is protective, presumably via removal of damaged mitochondria. While we also observe increased expression and mitochondrial localization of the mitophagy mediator PARKIN, loss of PARKIN does not worsen the cardiomyopathy seen in R120G CRYAB transgenic mice, unlike the findings seen in R120G CRYAB mice with ablation of one TRAF2 allele in cardiac myocytes. These observations indicate that TRAF2 mediates a beneficial upregulation of mitophagy in response to proteotoxic stress, in a PARKIN-independent manner. Furthermore, TRAF2 gain of function experiments demonstrate that stimulating mitophagy can be a translational strategy to remove protein aggregates, improve mitochondrial structure and function, and delay the development of cardiomyopathy. In this regard, our prior work indicates that AAV9-mediated transduction of TRAF2 to cardiac myocytes induces modest TRAF2 overexpression in the mouse heart (∼2.5 fold, as we have reported^27^ and as shown in Figure 7A) and stimulates mitophagy without an adverse effect on cardiac structure and function. Similarly, overexpression of PARKIN in cardiac myocytes in vivo did not adversely affect resting cardiac structure and function in absence of another stress,^46, 52, 53^ suggesting that modest upregulation of mitophagy (<2 fold with TRAF2^27^) is well tolerated in the murine myocardium.

The dramatic increase in myocardial TRAF2 abundance in mice expressing the R120G CRYAB mutant protein suggests a transcriptional response to coordinately upregulate mitophagy. Indeed, MiT/TFE family of transcription factors get activated upon induction of mitophagy and are required to sustain the process through transcriptional induction of the autophagy-lysosome machinery.^54^ TRAF2 is not a previously described target for TFEB suggesting alternate transcriptional responses may also coordinately drive mitophagy, which requires exploration in future studies. Interestingly, we have found that activation of TFEB, the canonical MiT/TFE family member, mediates the beneficial effects of intermittent fasting in mice expressing the R120G-CRYAB mutant protein.^16^ On a cautionary note, our collaborative studies point to potential toxicity with strategies for global stimulation of autophagy such as with sustained cardiac-myocyte TFEB overexpression (available as a preprint)^55^; and the potential for autophagic cell death or worsening cardiomyopathy from excess stimulation of autophagy.^56, 57^ Therefore strategies to target specific autophagy pathways^58^ such as mitophagy to remove damaged mitochondria or aggrephagy to remove protein aggregates may be preferred for minimizing off-target effects of global autophagy activation.

It is also intriguing that multiple mitophagy pathways have been described in cardiac myocytes to remove and degrade mitochondria, pointing to the critical need for maintaining normal mitochondria in cardiac myocytes, in the face of potentially high energetic costs involved in these long-lived and mostly irreplaceable cells. For example, in addition to the canonical mitophagy pathways discussed above, mitochondria can also be degraded via an alternative rab9-dependent pathway,^59^ via a rab5-dependent endosomal pathway,^60^ or released in exosomes to be phagocytosed and degraded within macrophage lysosomes.^61^ The existence of multiple mitophagy pathways points to the myriad critical roles for these organelles in cardiac myocyte biology. Our findings add to this understanding by uncovering another important function for mitochondria in cardiac physiology and stress response by importing and degrading cytosolic aggregate proteins. Notably, mitochondrial transplantation therapies are being pursued to ameliorate cardiac pathology in patients with mitochondrial DNA mutations, as well as the broader application of this technique to treat metabolic abnormalities in heart failure in on the horizon.^62^ The potential for mitochondrial therapies to improve sarcomere structure and function is an exciting possibility that merits experimental evaluation.

## Methods

### Reagents

We crossed mice homozygous for floxed *Traf*2 alleles (*Traf2* fl/fl)^63^ (which permits Cre-mediated excision of exon 3 and introduces a frame shift resulting in loss of TRAF2 protein; generously provided by Dr. Robert Brink, Garvan Institute of Medical Research, Australia) with mice carrying the *Myh6* promoter driven Mer-Cre-Mer (MCM) transgene (generous gift from Jeffery D. Molkentin, Ph.D., Cincinnati Childeren’s Hospital, Cincinnati, OH).^64^ Transgenic mice with cardiomyocyte-specific expression of R120G mutant of human CRYAB (namely the *Myh6*-CRYAB-R120G transgenic mice) were described previously.^37^ Mice were fed tamoxifen chow (ENVIGO, cat#TD.130857), as described for individual experiments. MitoQC reporter strain was previously described ^44^ and generously provided by Dr. Ian Ganley at University of Dundee, U.K. *Park2*-null mice were obtained from JAX (Strain number 006582). All mice were maintained on a C57BL/6J background and littermates were studied as controls. Mice of both sexes were studied. No significant differences were observed between sexes for the primary phenotype, whereby data for both sexes were combined for presentation. Mouse studies were randomized, and observers blinded. All animal studies were approved by the Institutional Animal Care and Use Committee (IACUC) at Washington University School of Medicine.

### Echocardiography

2D-directed M-mode echocardiography was performed as we have previously described ^16^.

### Studies with adeno-associated viral vectors

Adeno-associated virus serotype 9 (AAV9) particles coding for TRAF2 or GFP (which we have described previously^27^) driven by the cardiac troponin T promoter for conferring cardiac myocyte selective expression^65^, were generated by the Washington University Hope Center Viral Vectors Core; using AAV backbone constructs generously provided by Dr. Brent French at University of Virginia, Charlottesville, VA.

### Neonatal mouse cardiac myocyte isolation

Neonatal mouse cardiac myocyte isolation was performed from *Traf2* fl/fl mice using a modification of the technique we have described with the Worthington Neonatal Cardiomyocyte Isolation System (CAT# LK003300).^16^ Hearts were harvested from one-day old neonatal mice, and were subjected to trypsin digestion in a final concentration of 50 μg/ml in HBSS for 16-18 hours at 4°C after removal of the atria. Collagenase digestion (type II collagenase; 300 U/ml; Worthington) was conducted at 37°C for 45 min. Cardiomyocytes were seeded on collagen-coated four well chamber slides (Laboratory Tek) at a density of 10^5^ cells per square cm. On the 2nd day, the culture medium was changed to the Rat Cardiomyocyte Culture Medium (Cell applications INC, CAT#R313-500) for at 3-5 days prior to staining.

### Biochemical subcellular fractionation

Mitochondria-enriched and cytosolic fractions were prepared from hearts and cells following the protocols we have previously described.^27^

### Mitochondrial import assay

HEK293 cells were transfected with pcDNA3.1-CMV-GFP-CryAB or pcDNA3.1-CMV-GFP-CryABR120G mutant plasmid using Lipofectamine 3000 (Invitrogen, L3000–008) for 24 h according to the manufacturer’s instructions. Cytosolic fractions isolated from the GFP-CryAB or CryABR120G mutation expressing cells were admixed with mitochondria isolated from the untransfected cells, and incubated at room temperature for 1 h; and mitochondria were isolated via centrifugation, as previously described.^66^ To examine whether the imported GFP-CryAB or GFP-CryABR120G mutant proteins localize within the mitochondrial matrix, mitochondria harvested as above were further treated with 5 mg/ml of Digitonin (Abcam, ab141501) as described,^42^ for 10 min at room temperature prior to immunoblot analysis. Digitonin treatment at this concentration strips the outer mitochondrial membranes to generates mitoplasts, based on the differences in the solubility of the outer and inner mitochondrial membranes to this detergent.^67^ Mitochondria were pre-treated with 4mM CCCP for 5 minutes prior to incubation with cytosol from CRYAB R120G expressing HEK293 cells. To knockdown HSC70, HEK293 cells were treated with esiRNA targeting human HSC70 or firefly luciferase as control for 72 hours prior to the experiment (Sigma, human HSC70 ehu115141, firefly luciferase ehufluc, MISSION® siRNA Transfection Reagent, s1452).

### Immunofluorescence Analysis

Paraffin-embedded heart sections (10 µm thick) were subjected to heat-induced epitope retrieval, followed by blocking, and incubated overnight with primary antibody and subjected to the protocol we have previously described.^16^ After serial washes, samples were stained with indicated antibodies and mounted with fluorescent 4’,6-diamidino-2-phenylindole mounting medium (Vector Labs, H-1200). Confocal imaging was performed on a Zeiss confocal LSM-700 laser scanning confocal microscope using 639 Zeiss Plan-Neofluar 40/1.3 and 63/1.4 oil immersion objectives, and images were acquired using Zen 2010 software. The following primary antibodies were used for immunofluorescence imaging: p62 (Abcam, ab5416); DESMIN (Santa Cruz, SC7559), COXIV (Cell Signaling, 11967); Ubiquitin (Abcam, AB134953)

### Super resolution Microscopy

Super resolution microscopy of mouse heart sections was performed with the Zeiss LSM 880 Confocal with Airy scan at the Washington University Center for Cellular Imaging. IHC samples from *Traf2* floxed mice and *Traf2*-icKO mice labeled mitochondria with COXIV-FP (ex 600, em 650) was detected with the 633 laser, P62-GFP (ex395, em 509) was detected with the 488 laser, and Ubiquitin-RFP (ex 558, em 583) was detected with the 561 laser. Images were captured in z-stack using the 60x objective. Images were deconvolved using AiryScan processing to achieve 120 nm resolution.

### Assessment of mitophagy

Mitophagy was assessed in adult 20 week-old Myh6-R120G CRYAB transgenic mice carrying the mitoQC reporter allele as previous described.^27, 44^ Mitophagy was scored by counting mCherry-only puncta per field.

### Assessment of myocardial histology and ultrastructure

Staining for hematoxylin and eosin, and Masson’s Trichrome, was performed on formalin fixed cardiac tissue, and transmission electron microscopy was performed on glutaraldehyde fixed cardiac tissues, as previously described.^27^

### Assessment of proteasome activity

Proteasome activity examined by a commercially available fluorometric assay kit (Proteasome Activity Assay Kit, Abcam, ab107921) following manufacturer’s instruction.

### Immunoblotting

Immunoblotting was performed as previously described 30. Specific antibodies employed are as follows: p62 (Abcam, ab5416); TRAF2 (Abcam, ab126758); TOMM20 (Sigma, WH0009804M1); VDAC (Cell Signaling, 4661S); PARKIN (Cell Signaling, 2132); GAPDH (Abcam, ab22555); GFP (Abcam, AB290); LONP1 (Proteintech, 15440-1-AP); CLPP (Proteintech, 15698-1-AP); CRYAB (Enzo, ADISPA223F); DESMIN (Santa Cruz, SC7559), HTRA2 (Proteintech, 15775-1-AP); COXIV (Cell Signaling, 11967); Ubiquitin (Enzo, BML-PW8805-0500); Ubiquitin (Abcam, AB134953); SOD2 (Cell Signaling, 13141); HSC70 (Abnova, mab6636).

### Induced pluripotent stem cell derived cardiac myocytes

Induced pluripotent stem cells with mKate-tagged α-actinin were previously described.^40^ These cells were targeted with CRISPR-Cas9 approach at the Genome Editing and Stem Cell Center at Washington University School of Medicine to generate homozygous knock in lines to change codon for expression of arginine to glycine at position 120. The mutation was validated by next generation sequencing and resulting R120G mutant iPSC and isogenic controls were cultured in MTesR1 media before being transitioned to Essential 8 media (E8). During passaging and differentiation, iPSC were passaged as single cells using Gentle Dissociation Reagent (PBS with 1.8g/L NaCl and 0.5mM EDTA)^68^ and replated into media supplemented with 10μM Y27632 (Biogems-1293823) onto plates cultured with Geltrex (Life Technologies A1413302) at a concentration of 18.75μg/cm^2^. For differentiation, iPSC were subjected to timed control over Wnt signaling using small molecules.^69^ Briefly, iPSC were seeded at a density of 40k cells/cm^2^ onto Geltrex (37.5μg/cm^2^) and expanded for 3 days until confluent in E8. Next, media was changed to RPMI1640 supplemented with 150μg/mL ascorbic acid and 2% B27 without insulin (RPMI-I) containing 6μM CHIR99021 (Biogems-2520691, differentiation day 0). After 2 days, media was changed to RPMI-I containing 5μM IWP2. After 2 additional days, media changed to RPMI1640 supplemented with 2% B27 (RPMI-C, Biogems-6866167). Thereafter, cultured were fed every 2 days. Beating cardiomyocytes were typically observed by differentiation day 8. On differentiation day 14, beating cardiomyocyte monolayer were gently dissociated using TrypLE Select Enzyme (10X), Gibco A1217701. The single cells were then replated onto Geltrex-coated tissue culture plastic (37.5μg/cm^2^) at a density of 300,000 cells/cm^2^ and cultured in RPMI-C media supplemented with 20% fetal bovine serum (FBS) and 10μM Y27632 for 24hours. At day 15, the media was changed to a lactate-based metabolic selection media (RPMI1640 without glucose supplemented with 4mM lactate, 1% Non-Essential Amino Acids, and 1% Glutamax). On days 18 and 21, the media was replaced with fresh metabolic selection media. After 10 days culture in metabolic selection media, from day 25 to 29, the media was gradually changed back to RPMI-C for post-selection recovery.

### Assessment of mitochondrial function

High resolution oximetry was performed using the Oroboros Oxygraph 2k. A 6 well dish was prepared for incubating the heart and preparing it for the experiment. Cardiac tissue separated into thin fibers of ∼0.5-1.0 mg weight was used for the experiment. For each individual sample, 2 mL BIOPS solution was added to one well and 2 mL MirO5 (no additives) was added to another well. Then, 5 mg/ml Saponin solution was prepared in milliQ H2O. Dissected desired tissue from animals were placed on ice in 1 mL of BIOPS solution until all tissues are collected. 20 µL of Saponin solution was added to BIOPS well in the 6-well dish, on ice while rocking in the cold room for 20 minutes. While the tissue is permeabilizing, MirO5Cr solution was prepared by adding 15 mg of creatine powder to 5 mL of MirO5 and 5 µL of 10mM Blebbistatin stock to the MirO5 containing creatine. Pyruvate was prepared by adding 200 µL of H_2_O to 44 mg of pyruvate powder. When permeabilization is complete, fiber bundles were transferred to well containing MirO5 and incubated for 15 minutes. After MirO5 incubation is complete, the fibers were blotted dry and weighted for use. 2.1 mL of prepared MirO5 solution was used in the chamber wells where cardiac fibers were placed. ∼400 pmol of O_2_ was injected into the chamber and was air sealed. Then mitochondrial respiration was measured using the following substrates sequentially: a) 2.5 µL 800mM Malate, b) 10 µL 2M Glutamate and 5 µL of Pyruvate, c) 20 µL of 500mM ADP, d) 20 µL of 1M Succinate, e) 5 µL of 4mM Cytochrome C, f) 3 x 1 µL 1mM FCCP, g) 1 µL of 1 mM Rotenone. The next substrate was added only when the rate of O_2_ flux has leveled off after adding the previous substrate. Raw data were analyzed using the Oroboros DatLab software version 7.4.0.4. O_2_ flux was calculated by normalizing the weight of the cardiac tissue per mg.

### Assessment of aggregate abundance

To quantitate DESMIN aggregates in tissue sections, we utilized the following scoring system. Cardiac myocytes without aggregates were scored as 0 and cardiac myocytes with at least one DESMIN+ aggregate was scored as 1. The number of cardiac myocytes with aggregates were then divided by the total number of cardiac myocytes in the field of view. The scores were taken for five images for each heart and averaged. Image acquisition and quantitation was done by different operators with scoring done in blindly.

### Statistical analysis

Data are presented as mean ± standard error of the mean (SEM). All measurements were obtained on distinct biological replicates. Statistics were performed in Prism Version 9.1.2 (GraphPad Software Inc.). Data were tested for assumptions of normality with Shapiro-Wilk normality test. Statistical significance of differences was calculated via unpaired 2-tailed Student’s t test for 2 group comparisons, or one-way analysis of variance (ANOVA) for assessing differences across multiple groups followed by post-hoc testing (Tukey’s) to evaluate pairwise differences. A non-parametric test was employed if data were not normally distributed as indicated in the figure legend. A log-rank (Mantel-Cox) test was used to assess statistical significance in survival studies. Graphs containing error bars show means ± SEM with a P value < 0.05 considered statistically significant. The datasets from the current study are available from the corresponding author on reasonable request.

### Sources of funding

A.D. was supported by grants from the National Institutes of Health (HL107594, HL143431 and NS094692) and the Department of Veterans Affairs (I01BX004235, I01BX005065, I01BX005981). D.R.R. is supported from grants from the National Institutes of Health (T32 HL007081 and 1K08HL163469). This study was also supported by a Shared Instrumentation Grant from the NIH (S10 OD028597) to A.K.

## Author contributions

D.R.R., M.I., C.Z., J.T.M., H.N., M.K., X.G., J.N., Y.K.G.K., A.K. A.M. and X.M. performed experiments, acquired, and analyzed the data; N.H. provided critical reagents, and K.M. and N.H. interpreted the data and revised the manuscript; D.R.R., X.M. and A.D. conceived the experiments, analyzed the data, supervised the work, and drafted the manuscript.

## Supplementary Figures

**Figure S1.**
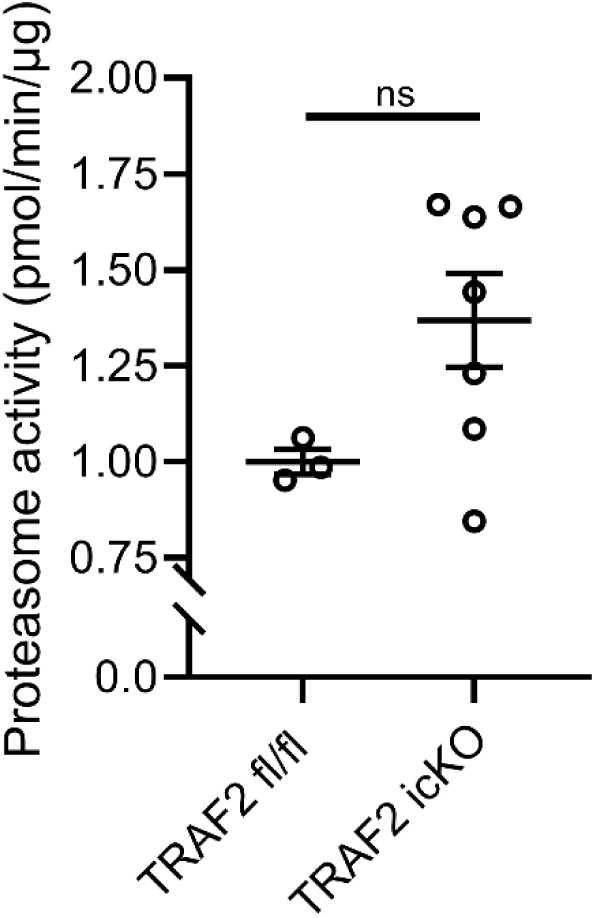
Cardiac myocyte deletion of TRAF2 does not decrease cardiac proteasome activity. Proteasome activity assessed in heart tissue from TRAF2-icKO mice versus TRAF2 fl/fl controls, modeled as in Figure 1A. Graph shows mean +/-SEM. No statistically significant difference was detected by t-test.

**Figure S2.**
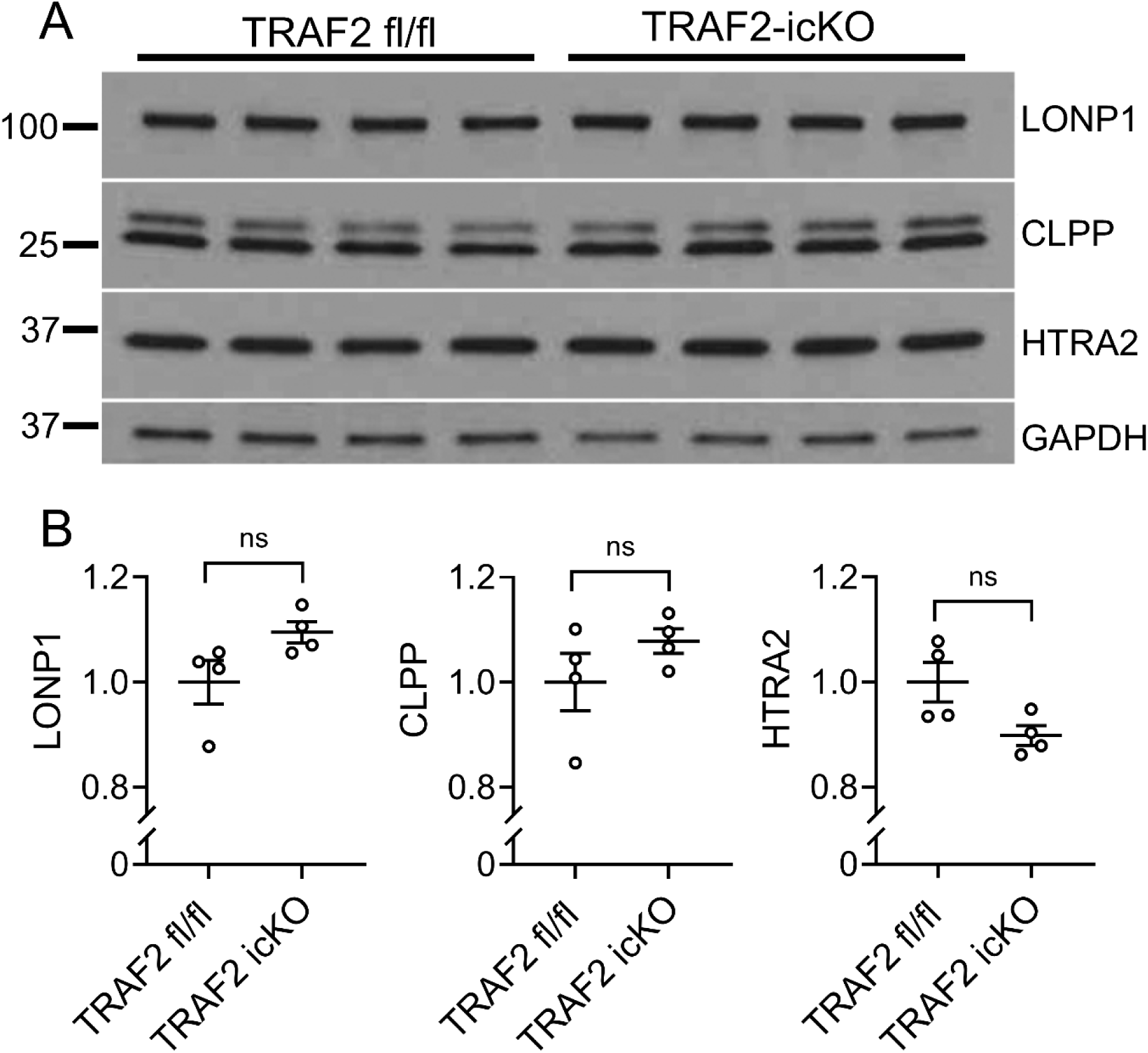
TRAF2 deletion does not alter mitochondrial protease levels. **A** Immunoblotting for mitochondrial proteases LONP1, CLPP, and HTRA2 in TRAF2-icKO and TRAF2 fl/fl hearts, modeled as in Figure 1A. **B** Quantitation of LONP1, CLPP, and HTRA2 levels from A. Levels are normalized to GAPDH as the loading control. No statistically significant differences were detected between groups by t-test.

**Figure S3.**
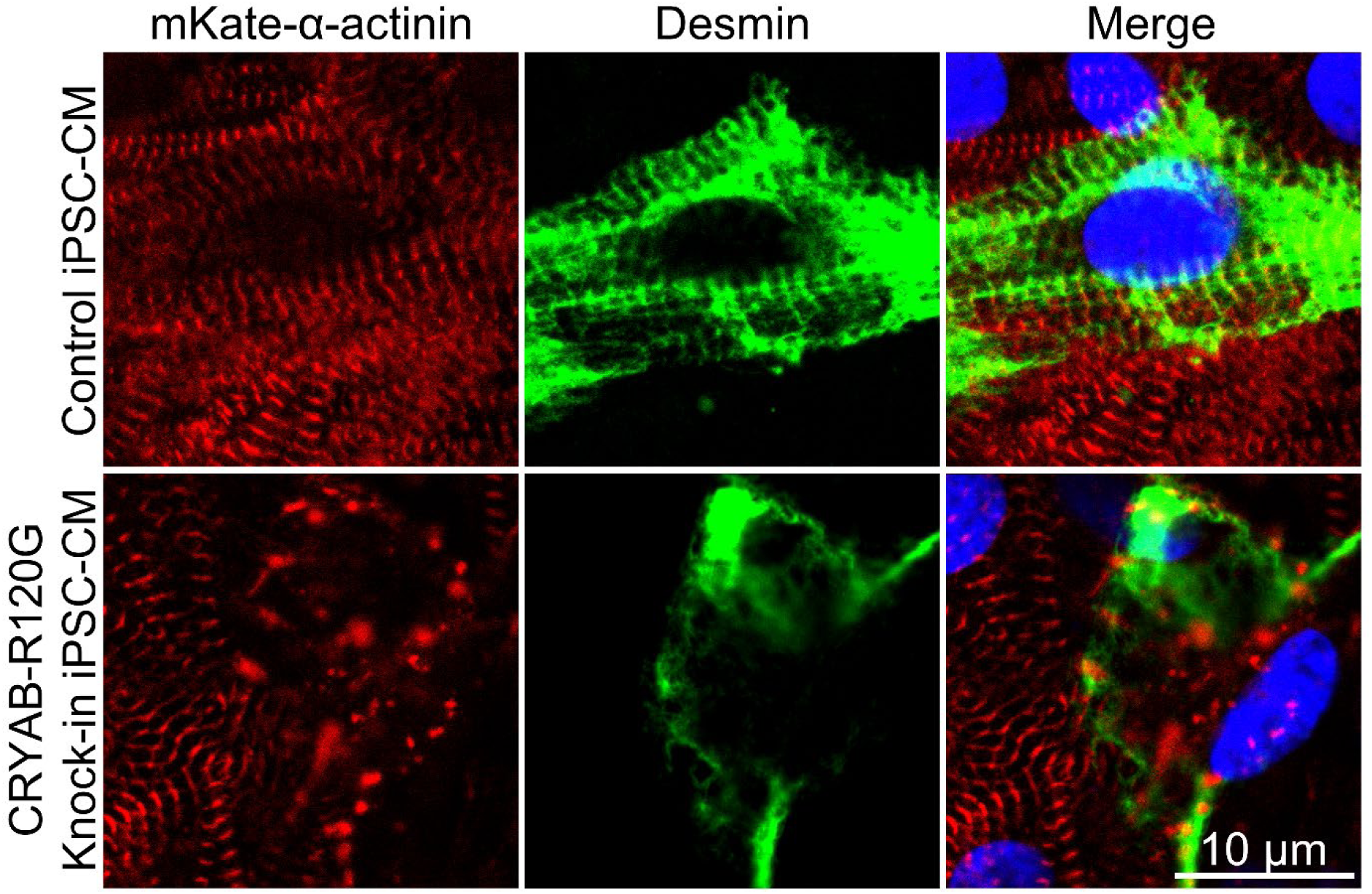
DESMIN aggregates are observed in R120G knock-in iPS cell derived cardiac myocytes. Representative images from induced pluripotent stem cell (iPSC)-derived cardiac myocytes (cultured in aggregates for 30 days) from iPSC lines homozygous for knock-in of R120G mutation and isogenic controls expressing mKate-tagged α-actinin and stained with antibody against DESMIN. Images demonstrate localization of DESMIN and α-actinin aggregates in R120G knock-in iPSC-derived cardiac myocytes (bottom panel), as compared with localization of these proteins in a sarcomeric pattern in control iPSC-derived cardiac myocytes (top panel). DAPI stained nuclei are blue.

**Figure S4.**
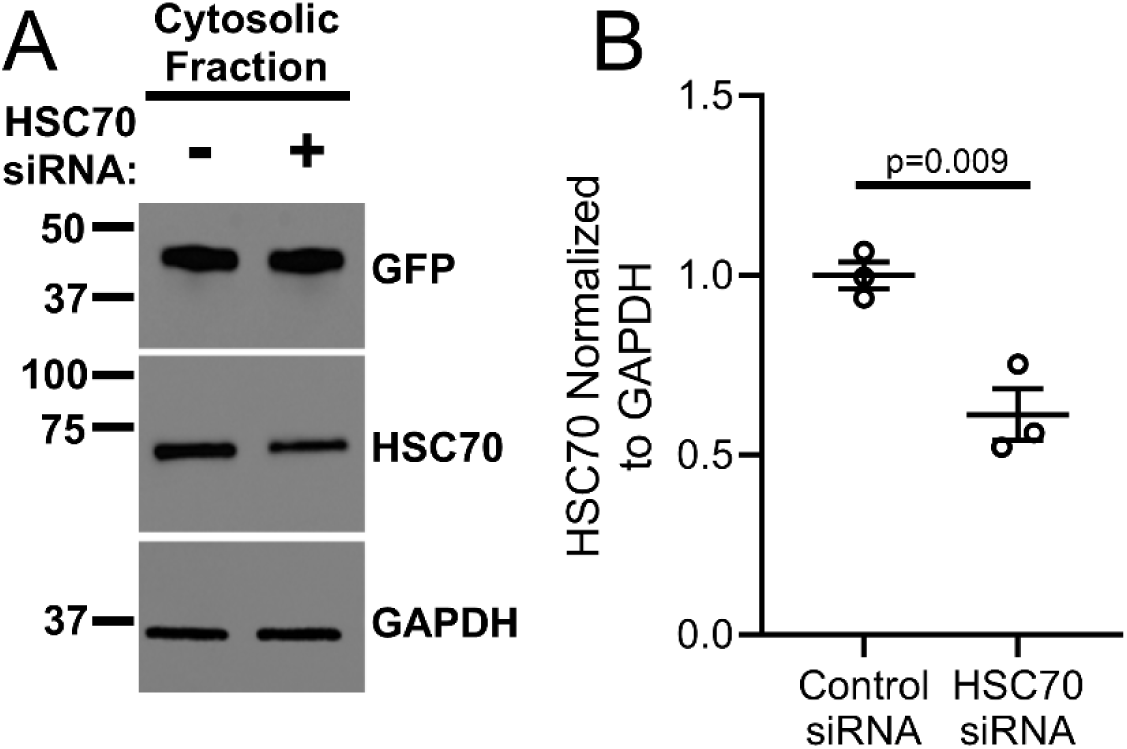
siRNA transfection results in knockdown of HSC70 in HEK293 cells. **A, B** Representative immunoblot (A) depicting expression of HSC70 and GFP in GFP-tagged R120G CRYAB transfected with siRNA targeting HSC70 (depicted as ‘+’) or scrambled control (depicted as ‘-‘) in HEK293 cells with quantitation of HSC70 abundance (B, expressed as fold change over control). GAPDH is employed as loading control.

**Figure S5.**
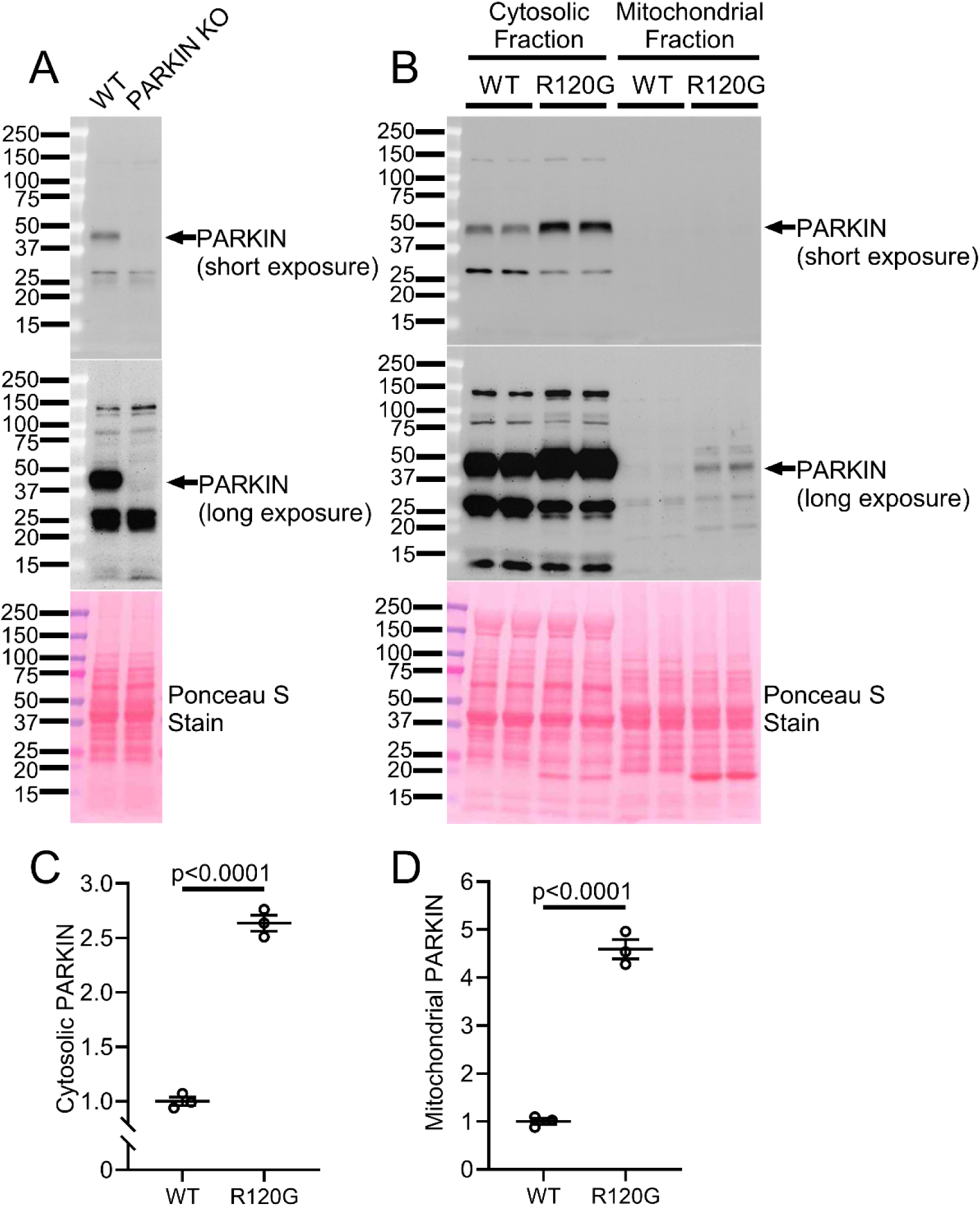
PARKIN expression and mitochondrial localization is increased in CRYAB-R120G hearts. **A** Immunoblotting in wild-type (WT) and Park2 null (PARKIN KO) heart samples demonstrating specificity of the PARKIN antibody. Arrow indicates PARKIN-specific band. **B** Immunoblotting on cytosolic and mitochondrial fractions from WT and CRYAB-R120G hearts at 20-24 weeks of age. PARKIN blots show short and long exposures. Cytosolic and mitochondrial samples are from the same fractionations shown in Figure 5E; see this panel for cytosolic (GAPDH) and mitochondrial (COXIV, VDAC) markers to demonstrate successful separation of mitochondrial and cytosolic fractions in these samples. **C-D** Quantitation of PARKIN levels in cytosolic and mitochondrial fractions in immunoblotting in panel b. Samples are normalized to total protein as shown by Ponceau S staining. *P* values shown are by t-test.

**Figure S6.**
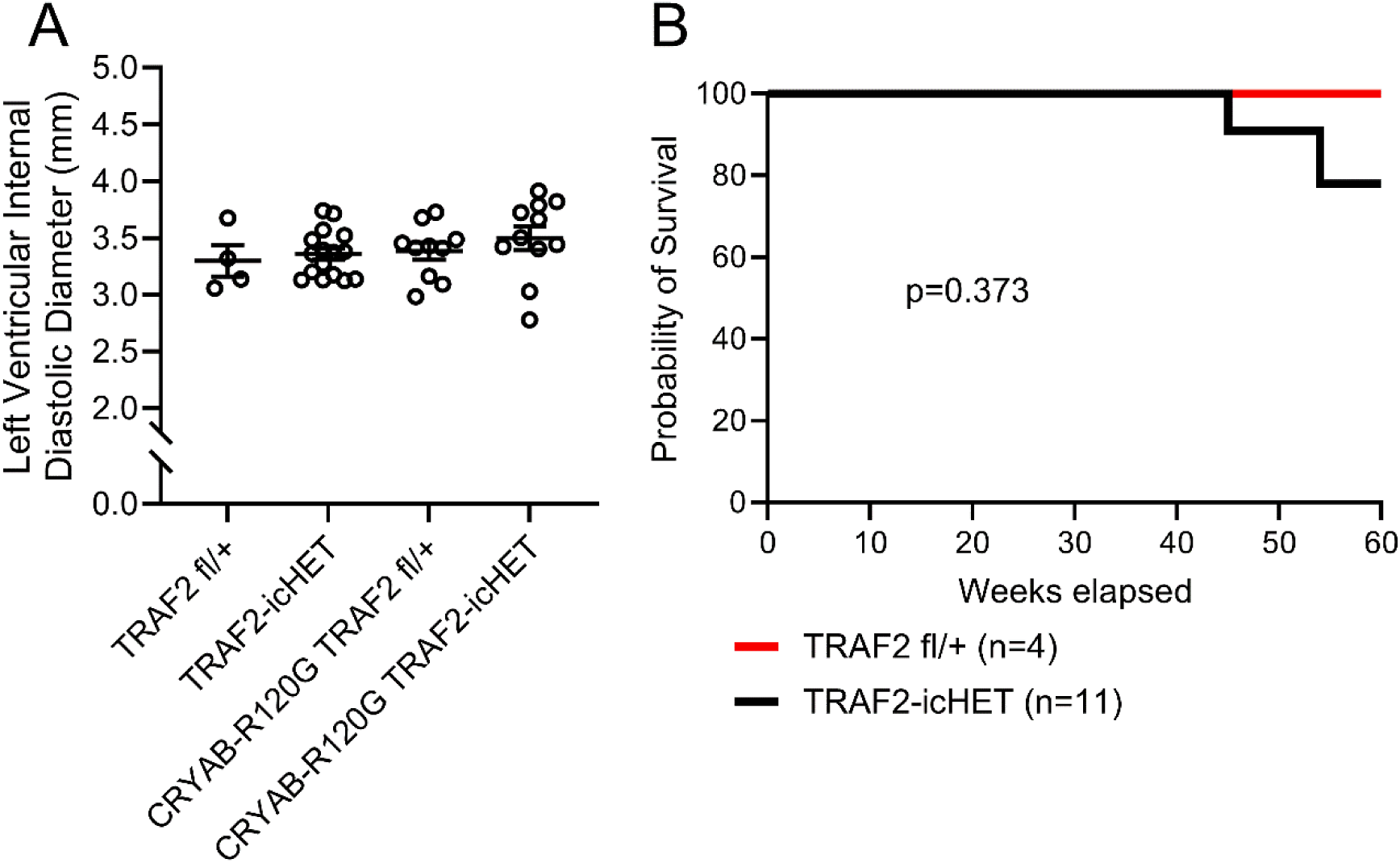
Reduction in cardiac myocyte TRAF2 does not affect left ventricular dimensions in Myh6-R120G CRYAB transgenic mice or alter mortality in wild type mice. **A** Left ventricular internal diastolic diameter by M-mode echocardiography in 20-week-old CRYAB-R120G mice with cardiac myocyte-specific inducible deletion of one *Traf2* allele (termed R120G TRAF2-icHET; see Figure 6A for experimental schematic). These mice were compared to R120G TRAF2 fl/+ mice without the MerCreMer transgene, and to TRAF2 fl/+ and TRAF2-icHET control mice without CRYAB-R120G. Statistical comparisons were performed by one-way ANOVA followed by Tukey’s test for multiple comparison testing between groups. There were no statistically significant differences between groups in panel A. **B** Kaplan-Meier survival analysis of TRAF2-icHET mice and TRAF2 fl/+ controls. No statistically significant differences were detected by Mantel-Cox log-rank testing.

**Figure S7.**
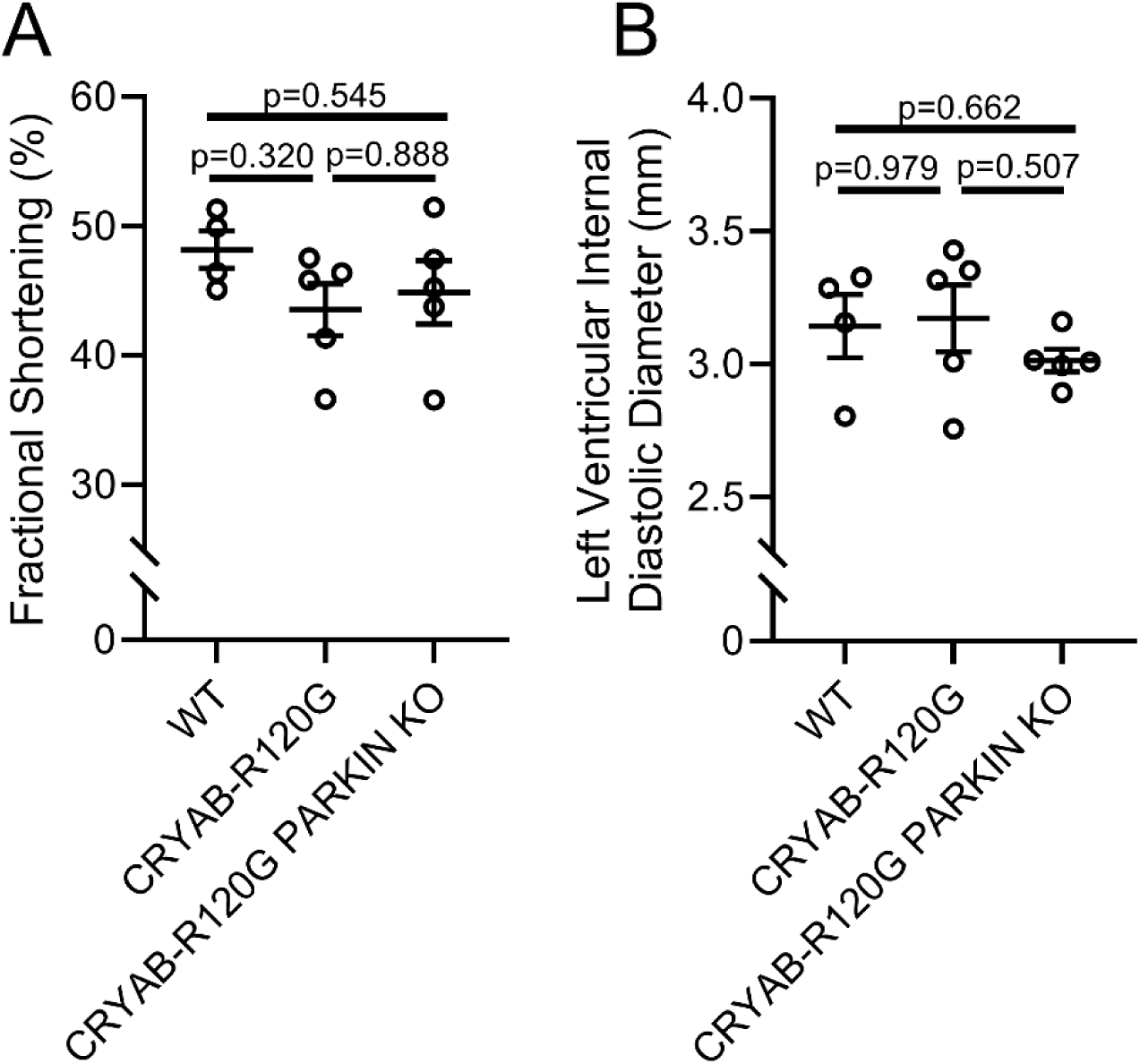
PARKIN ablation in Myh6-R120G CRYAB transgenic mice does not affect left ventricular structure and function. **A, B** Left ventricular endocardial fractional shortening (A) and left ventricular internal diastolic diameter (B) by M-mode echocardiography in 20-week-old CRYAB-R120G mice with and without concomitant *Park2* null alleles (as homozygous, PARKIN KO) and wild type as control. No statistically significant differences were detected by one-way ANOVA followed by Tukey’s test for multiple comparison testing between groups.

**Figure S8.**
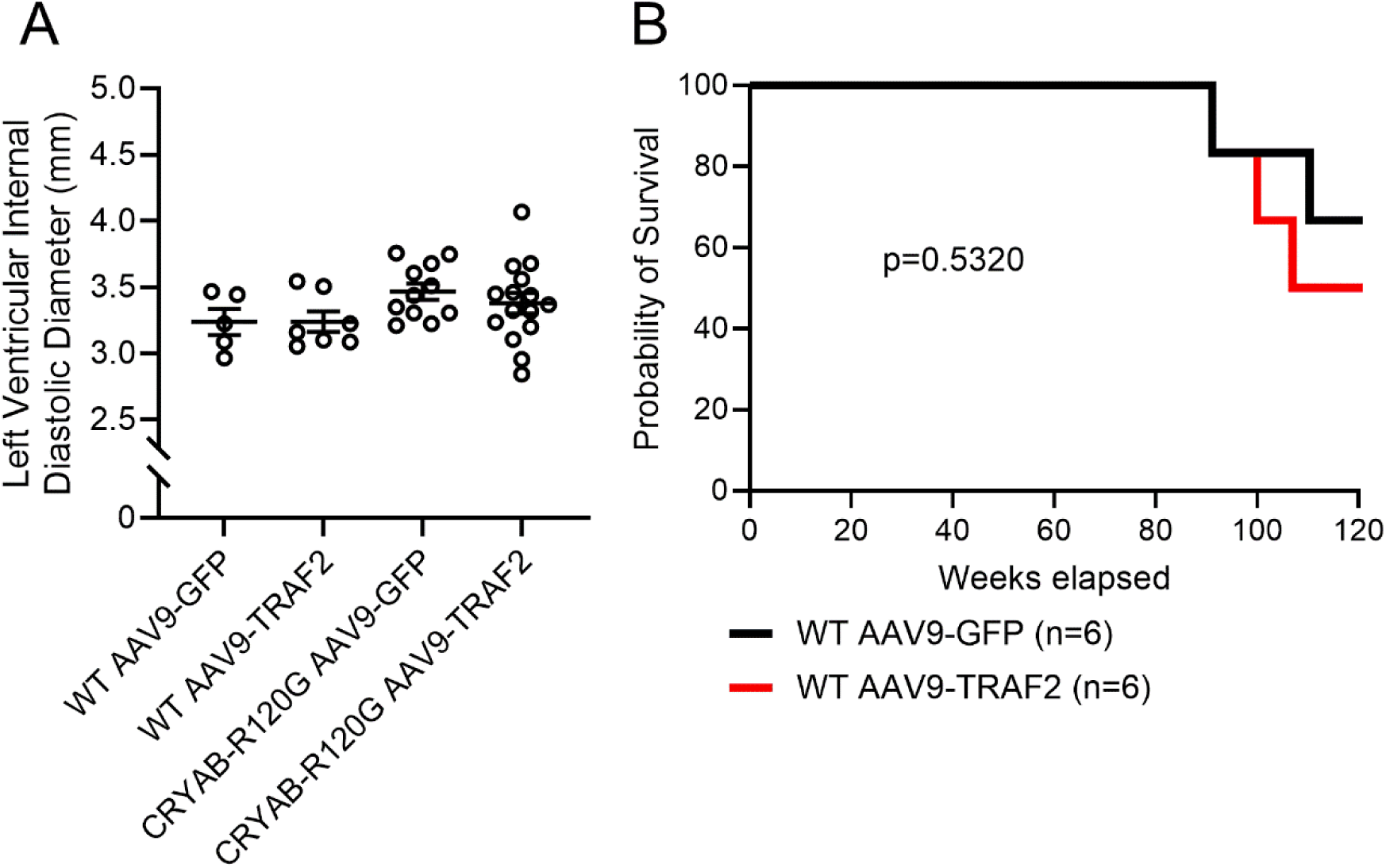
AAV9-mediated transduction of TRAF2 does not affect left ventricular dimension in Myh6-R120G CRYAB transgenic mice or alter mortality in wild type mice. **A** Left ventricular internal diastolic diameter by M-mode echocardiography in 20-week-old CRYAB-R120G mice and wild type mice with AAV9-cardiac Troponin T promoter driven transduction of TRAF2 or GFP at 8 weeks of age (See Figure 8A for experimental schematic). Statistical comparisons were performed by one-way ANOVA followed by Tukey’s test for multiple comparison testing between groups. There were no statistically significant differences between groups in panel A. **B** Kaplan-Meier survival analysis of wild type mice with AAV9-cardiac Troponin T promoter driven transduction of TRAF2 or GFP at 8 weeks of age. No statistically significant differences were detected by Mantel-Cox log-rank testing.

**Figure S9.**
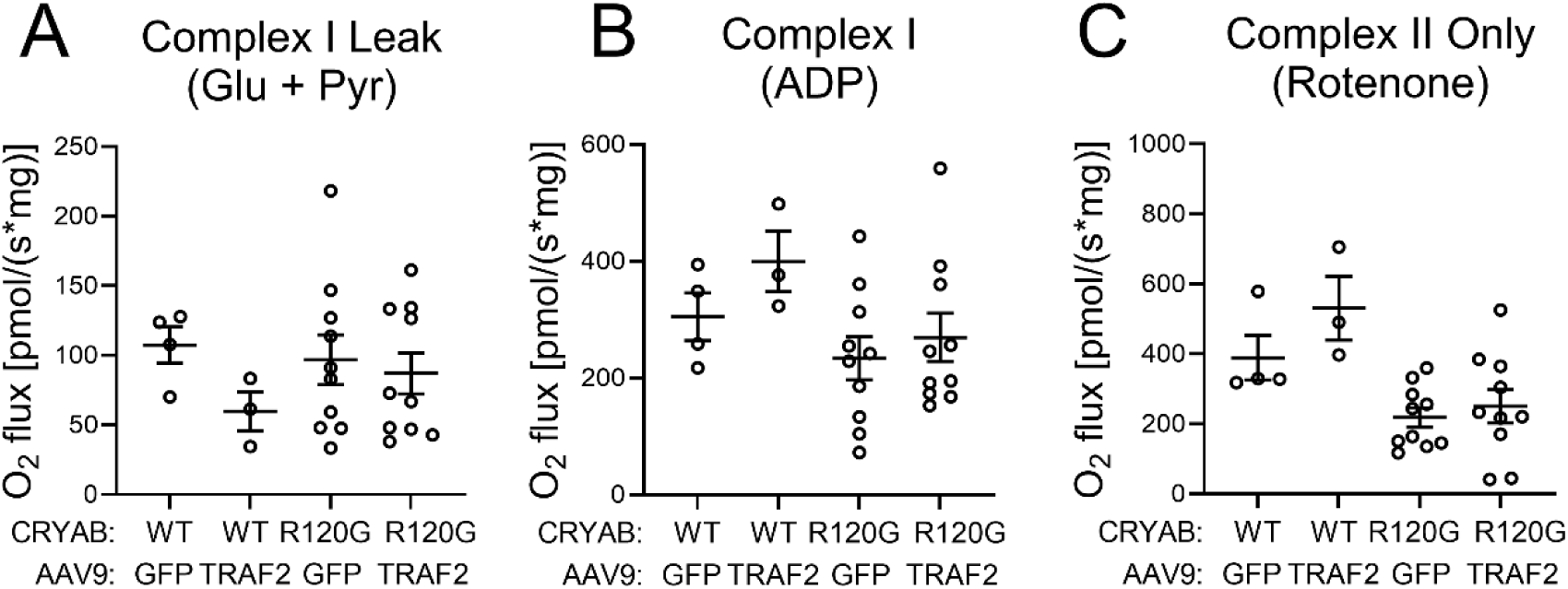
High resolution respirometry in Myh6-R120G CRYAB transgenic mice transduced via AAV9-GFP or AAV9-TRAF2. **A-C.** High-resolution respirometry was performed to measure oxygen consumption [volume-specific oxygen flux (JO2)] in isolated, permeabilized cardiac fiber bundles from mice modeled as in Figure 8a. Oxygen consumption was assessed in the presence of (A) glutamate and pyruvate (to assess complex I leak), (B) ADP (to assess complex I activity), and (C) rotenone (to assess complex II activity). See Methods section for complete description of experimental methods. Statistical comparisons were performed by one-way ANOVA followed by Tukey’s test for multiple comparison testing between groups in A, and by Kruskal-Wallis test followed by Dunn’s test for multiple comparison testing between groups in B and C. Statistically significant differences between groups are shown in panels; all other comparisons were p>0.05.

